# Norepinephrine regulates Ca^2+^ signals and fate of oligodendrocyte progenitor cells in the cortex

**DOI:** 10.1101/2022.08.31.505555

**Authors:** Frederic Fiore, Ram R. Dereddi, Khaleel Alhalaseh, Ilknur Coban, Ali Harb, Amit Agarwal

## Abstract

Oligodendrocyte precursor cells (OPCs) represent the most abundant group of proliferating cells in the adult central nervous system. OPCs serve as progenitors for oligodendrocyte (OLs) throughout the life, and contribute to developmental and adaptive myelination, and myelin repair during diseased state. OPCs make synaptic and extra-synaptic contacts with axons, and detect and respond to neuronal activity. How OPCs translate the information relayed by the neuronal activity into Ca^2+^ signals, which in turn influence their fate and survival, is less understood. We developed novel transgenic mouse lines expressing a cytosolic and membrane anchored variants of genetically encoded Ca^2+^ sensors (GCaMP6f or mGCaMP6s) in OPCs, performed 2-photon microscopy in the somatosensory cortex of the awake behaving mice, and simultaneously monitored intracellular Ca^2+^ signals and their cell-fate progression. We found Ca^2+^ signals in OPCs mainly occur within processes and confine to micrometer-size segments called Ca^2+^ microdomains. Microdomain Ca^2+^ signals enhanced in OPCs when mice engage in exploratory behavior. OPCs exhibit distinct Ca^2+^ signals while they proliferate to maintain their precursor pool or differentiate to generate new OL. When mice engaged in exploratory behavior, the cortical projections of noradrenergic neurons in locus coeruleus showed increased firing rate and norepinephrine release. Norepinephrine activated all three subtypes of alpha1 adrenergic receptor expressed by OPCs and evoked intracellular Ca^2+^ increase in OPCs. A chemogenetic activation of noradrenergic neurons, promoted differentiation of cortical OPCs into OL, and at the same time suppressed OPC proliferation rate. Hence, we uncovered that various cell types of oligodendrocyte lineage exhibits unique signatures of Ca^2+^ activity, which these cells might integrate for making their fate decisions, and norepinephrine signaling can be a potent regulator of OPC fate.

## Introduction

Oligodendrocytes (OLs) are myelinating cells in the central nervous system (CNS). They are post-mitotic cells and can survive for the entire lifespan of an organism^1, 2^. Unlike neurons and astrocytes, OLs are continually produced throughout adulthood^3^ from a pool of oligodendrocyte progenitor cells (OPCs) constituting 5-8% of all brain cells. Although myelination is thought to end in young adults, it remains incomplete in most brain areas. For example, about 20-40% of axons are unmyelinated in the corpus callosum^4^, and large axonal segments of excitatory and inhibitory neurons exhibit discontinuous myelination in the cortex^5^. This partial myelination in the CNS opens up a window for the formation of new myelin sheaths on unmyelinated axonal segments. Newly layered myelin extends metabolic support to neurons^6^, fine-tunes conduction velocity of individual axons and consequently influence neural circuit function^7, 8^. Hence, myelin remodeling is thought to be yet another form of long-term plasticity contributing to adaptive changes in brain function^9^.

OLs can remodel myelin by extending and retracting existing sheaths, or by adding entirely new internodes^10, 11^. The most common form of myelin remodeling originates from OPCs differentiating into new OLs, which then myelinate previously unmyelinated axonal segments^10^. Before OPCs differentiate into myelinating OLs, they integrate numerous cues in their local environment including neuronal activity and growth factors^12^. Rather than evolving unique signaling mechanisms, OPCs detect and respond to neuronal activity using the same pathways engaged by neurons during synaptic transmission. OPCs form direct synapses with unmyelinated axons^13, 14^ and express a wide variety of synaptic and extra synaptic neurotransmitter receptors^15–17^, thereby allowing them to engage in rapid communication. Axon-OPCs synaptic junctions are rapidly lost as OPCs begin to differentiate into OLs^16, 17^. This implies that, within the oligodendrocyte lineage, OPCs are uniquely endowed with the ability to sense and respond rapidly to electrical and chemical signals emanating from axons^18^.

Neuromodulators including acetylcholine (ACh), norepinephrine (NE) and serotonin (5-HT) can influence brain-wide neuronal activity^19^. In addition to the classical neurotransmitters such as glutamate and GABA, which act through local synaptic activity, OPCs in the adult brain might integrate extra-synaptic inputs from neuromodulators to guide their behavior. Indeed, several transcriptomics studies indicate that distinct subtypes of oligodendrocyte lineage cells (OLCs) express a wide variety of these neuromodulator receptors^20, 21^ which can affect their proliferation and differentiation^22, 23^. Notably, NE is a key neuromodulator involved in crucial biological processes including arousal, attention, memory formation, sleep/wake cycles and stress management^24^. NE has also been shown to regulate the proliferation and differentiation of neural progenitors^25, 26^. Although NE imparts immense influence on the neuronal function, its effect on OPC fate and myelination is elusive.

Neuromodulators such as NE exert their influence on target cells through the activation of G-protein coupled receptors (GPCRs), and often involve Ca^2+^ as a second messenger. Ca^2+^ is a universal signaling ion, and intracellular changes in Ca^2+^ concentrations appear to modulate the fate of OPCs by regulating proliferation, differentiation and programmed cell death^8^. Recent single-cell RNA-sequencing studies on rodent brains have identified a dozen cell types constituting the oligodendrocyte lineage (e.g., OPCs, premyelinating OLs and OLs)^20^, each expressing a distinct repertoire of neurotransmitter and neuromodulator receptors which grants them a unique ability to interact with the neighboring cells. During development, OPCs can experience varying levels of local Ca^2+^ changes based on their location, level of synaptic and extra-synaptic connectivity and type of neurotransmitter receptor expressed, which in turn can influence their fate. In the larval zebrafish, it has been shown that, during development, the characteristics of Ca^2+^ signal experienced by OPCs can regulate their differentiation into OLs^27^ as well as the stability of newly formed myelin internodes^28, 29^. Although these studies highlight the significance of Ca^2+^ signaling in regulating developmental OL generation and myelination, little is known about the characteristics of Ca^2+^ activity or the pathways that regulate Ca^2+^ signals in OPCs in the mature mammalian brain.

Like neurons, the source, location and amplitude of Ca^2+^ signals in OPCs might encode information about their developmental stage and fate progression. Hence, to understand how Ca^2+^ signals regulate the fate of OPCs, we performed high-resolution imaging of intracellular Ca^2+^ transients across the entire oligodendrocyte lineage, monitored cell fate and chemogenetically manipulated neuronal activity. To simultaneously study the Ca^2+^ signals and map the fate of OPCs, we developed novel transgenic mouse lines expressing two variants of a genetically encoded Ca^2+^ sensors and a red fluorescent reporter in OPCs, and performed chronic 2-photon microscopy on awake behaving mice. We found that Ca^2+^ signals in OPCs mainly occur within processes, and that OPCs exhibit distinct Ca^2+^ signals while they proliferate or differentiate. We also found that NE release from neurons evokes intracellular Ca^2+^ increases in OPCs, and suppresses their proliferation and promote differentiation.

## Results

### Generation and characterization of dual-color transgenic mouse lines for simultaneous fate mapping and Ca^2+^ imaging in OPCs

Unlike neurons and astrocytes, OPCs *in vivo* are difficult to transfect using recombinant AAVs carrying fluorescent proteins, genetically encoded activity reporters, and optogenetic effectors. In addition, there is no small-size promoter sequence that can reliably and efficiently drive a transgene expression in OPCs. Hence, it has been difficult to record (e.g., Ca^2+^ signals) and exclusively modulate the activity of OPCs and their lineage cells *in vivo*. To simultaneously map the fate and study changes in the Ca^2+^ signals of OPCs while they proliferate or differentiate into oligodendrocytes (OLs), we took a transgenic approach and generated two distinct triple-transgenic mouse lines (Fig. 1 and Extended Data Fig. 1). To capture an entire spectrum of Ca^2+^ transients (fast and slow) in the soma as well in the processes of OPCs, we used a mouse line conditionally expressing a cytosolic variant of ultrafast genetically encoded green fluorescent Ca^2+^ sensor GCaMP6f^30^ *(TIGRE2-tTA-LSL- GCaMP6f*) (Fig. 1a). To generate the first triple-transgenic mouse line, we cross bred these GCaMP6f mice with widely used *NG2-CreER* BAC transgenic mice^31^ expressing tamoxifen inducible variant of Cre recombinase (CreER) in OPCs, and a Cre-reporter mouse line expressing cytosolic variant of a red fluorescent protein tdTomato^32^ *(Rosa26-LSL-tdTomato*) (Fig. 1a). At 3-4 weeks of age, we injected the resulting triple transgenic mice (*NG2- CreER*;*TIGRE2-tTA-LSL-GCaMP6f*;*Rosa26-LSL-tdTomato;* in short referred as *NG2- GC6f;tdT*) with tamoxifen to induce the expression of GCaMP6f and tdTomato reporters in OPCs (Fig. 1b). 2-3 weeks post-tamoxifen injection, we characterized the efficiency and specificity of GCaMP6f and tdTomato expression by performing immunohistochemical analysis in the somatosensory cortex (S1; barrel cortex) of coronal brain sections from *NG2- GC6f;tdT* mice using antibodies against tdTomato, eGFP (GCaMP6f is a GFP derivative) and several oligodendrocyte lineage cells (OLCs) specific markers (Fig. 1b-e). Our histological analysis revealed that majority of recombined cells co-expressed tdTomato and GCaMP6f (Fig. 1b). However, a small fraction of cells exclusively expressed tdTomato and had a pericyte-like morphology (Fig. 1b, arrowheads). In the S1 cortex, almost all tdTomato/GCaMP6f double positive cells co-labeled with a pan oligodendrocyte lineage cells marker Olig2 (Fig. 1c). Both tdTomato and GCaMP6f co-expressed at high levels by PDGFRα+ OPCs (Fig. 1d) and ASPA+ OLs (Fig. 1e), and revealed both the soma and the processes of OPCs and OLs. A high expression of tdTomato and GCaMP6f is key for tracking the fate transitions of OPCs based on changes in the cell morphology, and for recording Ca^2+^ transients in soma and the fine processes of OLCs, respectively.

**Figure 1.**
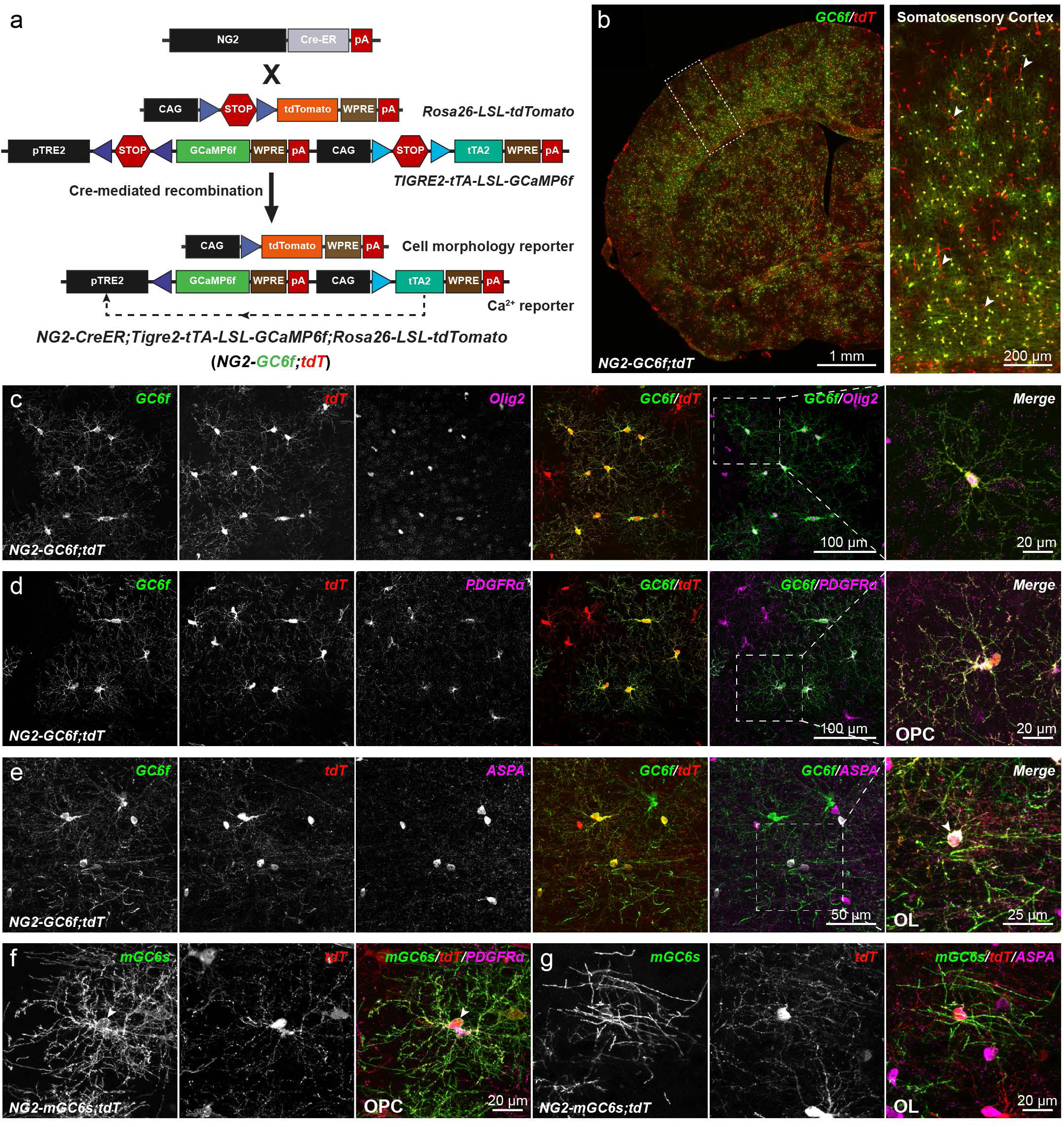
Conditional expression of GCaMP6f, mGCaMP6s and tdTomato in oligodendrocyte lineage cells. (**a**) Cartoon showing a triple transgenic strategy to express the ultrafast variant of GCaMP6 (GCaMP6f) and tdTomato in OPCs using *NG2-CreER;Rosa26-LSL-tdTomato;TIGRE2-tTA- LSL-GCaMP6f* (*NG2-GC6f/tdT*). (**b**) (left) Coronal brain hemi-section from a *NG2-GC6f/tdT* mouse stained for GCaMP6f (GC6f, green) and tdTomato (tdT, red). (right) Zoom-in image from the somatosensory cortex (boxed area shown in the left panel) shows oligodendrocyte lineage cells co-expressing GC6f and tdT. Arrowheads show recombined pericytes were mostly positive for tdT, but not for GC6f. Scale bars, 1 mm (left) and 200 µm (right). (**c**) Confocal microscopy images show double recombined (GC6f^+^/tdT^+^) cells express pan oligodendrocyte lineage marker Olig2. (right) High-magnification of the boxed area in **c**. Scale bars, 100 µm (left) and 20 µm (right). (**d**) Images showing a subset GC6f^+^/tdT^+^ cells were PDGFRα^+^ OPCs . (right) High-magnification image of GC6f^+^/tdT^+^ OPC in the boxed area in **d**. Scale bars, 100 µm (left) and 20 µm (right). (**e**) Images showing a subset of recombined GC6f^+^/tdT^+^ cells expressing ASPA, a marker for mature oligodendrocytes. (right) High-magnification of a GC6f^+^/tdT^+^ oligodendrocyte in the boxed area in **e**. The arrow highlights the cell body of a recombined mature oligodendrocyte. Scale bars, 50 µm (left) and 25 µm (right). (**f**) A recombined OPC from the triple transgenic mouse *NG2- CreER;Rosa26-LSL-tdTomato;ROSA-LSL-mGCaMP6s* (*NG2-mGC6s;tdT*) in the somatosensory cortex immunostained for PDGFRα, mGCaMP6s and tdT. The arrow highlights the membrane localization of mGC6s (green) in OPCs. Scale bar, 20 µm. (**g**) A recombined cell from *NG2-mGC6s;tdT* cortex expressing the mature OL marker ASPA. Scale bar, 20 µm.

Often, cytosolic reporter proteins do not reach fine processes of glial cells^33^ and axonal compartments of neurons^34^. Hence, to record Ca^2+^ transients in OPC processes and axonal projections, we generated a novel ROSA26 targeted knock-in mouse line which can conditionally express a membrane anchored variant of a sensitive genetically encoded green fluorescent Ca^2+^ sensor GCaMP6s (mGCaMP6s) in a cell-type specific manner (Extended Data Fig. 1). To generate the second triple-transgenic mouse line, we cross bred *NG2- CreER* and tdTomato reporter mice with our newly generated mouse line expressing mGCaMP6s *(Rosa26-LSL-mGCaMP6s*) (Extended Data Fig. 1a) to get *NG2-CreER*;*Rosa26- LSL-mGCaMP6s;Rosa26-LSL-tdTomato;* (*NG2-mGC6s;tdT*) mice. 3-4 weeks old mice were injected with tamoxifen to induce expression of tdTomato and mGCaMP6s in OPCs (Extended Data Fig. 1a). Similar to *NG2-GC6f;tdT* mice, our detailed histological analysis in the S1 cortex of *NG2-mGC6s;tdT* mice revealed that after Cre mediated recombination almost all cells double positive for tdTomato and mGCaMP6s co-labeled with pan oligodendrocyte lineage cell marker Olig2+ (Extended Data Fig. 1b-d). The majority of tdTomato+/mGCaMP6s+ cells were PDGFRα+ OPCs (Extended Data Fig. 1e) and a small subset of the double+ cells were ASPA+ OLs (Extended Data Fig. 1f). As expected, for a membrane anchored protein, mGCaMP6s only outlined the soma and didn’t fill the cytosol (as seen for tdTomato) of the OPCs (Fig. 1f) and OLs (Fig. 1g), but revealed the processes of OPCs and OLs in a great detail.

### Ca^2+^ signals in OPCs are restricted to Ca^2+^ microdomains and are regulated by neuronal activity

Very little is known about the characteristics of Ca^2+^ signals and the pathways activating these signals in OPCs *in vivo*. To study cell morphology and Ca^2+^ signals of cortical OPCs in mice freely engaging in explorative behavior, we adapted a high-precision mouse mobility tracking setup called Mobile HomeCage^35^ (MHC). MHC permitted head-fixation, a requirement for a high-resolution 2-photon microscopy, but otherwise allowed mice to “freely” explore their cage (Fig. 2a). To characterize the entire spectrum of fast and slow Ca^2+^ signals in the soma and processes of the OPCs, we used *NG2-GC6f;tdT* mice expressing a cytosolic ultrafast Ca^2+^ sensor GCaMP6f, which is capable of capturing Ca^2+^ transients in milliseconds range^36^. After tamoxifen induced expression of tdTomato and GCaMP6f in OPCs, we implanted a glass cranial window over S1 cortex, and trained mice over the course of a week to remain head-fixed for 60 minutes in the MHC environment (Fig. 2a, details in the Methods section). When mice reached adulthood (>9 weeks old), we performed a high-resolution dual color 2P-microscopy of OPCs at a 5 Hz image acquisition rate through the cranial window. We imaged simultaneously cell morphology (Fig. 2b, c) and Ca^2+^ signals in a single OPC (Fig. 2b, d). Since tdTomato and GCaMP6f signals were captured concomitantly, static tdTomato signals enabled a pseudo ‘ratiometric’ analysis of Ca^2+^ signals and the correction of mouse movement artifacts. We adapted our machine-learning based analysis software CaSCaDe^37^ to implement algorithms to automatically correct movement artifacts and analyze Ca^2+^ signals and mouse locomotion activity (see methods for details). We observed that Ca^2+^ transients in OPCs mostly occur in the processes and remain restricted to small segments – we termed such spatially confined micrometer size (4.64 ± 3.33 µm^2^) regions in OPCs as **Ca^2+^ microdomains** (CaMs) (Fig. 2d- f). Binarized Ca^2+^ signals traces from neighboring CaMs indicate that each CaM can exhibit changes in Ca^2+^ concentration independently, but can also occasionally be activated alongside several other CaMs (Fig. 2e: A-H; Fig. 2g), thereby suggesting that CaMs in OPCs can act as independent signaling units.

**Figure 2.**
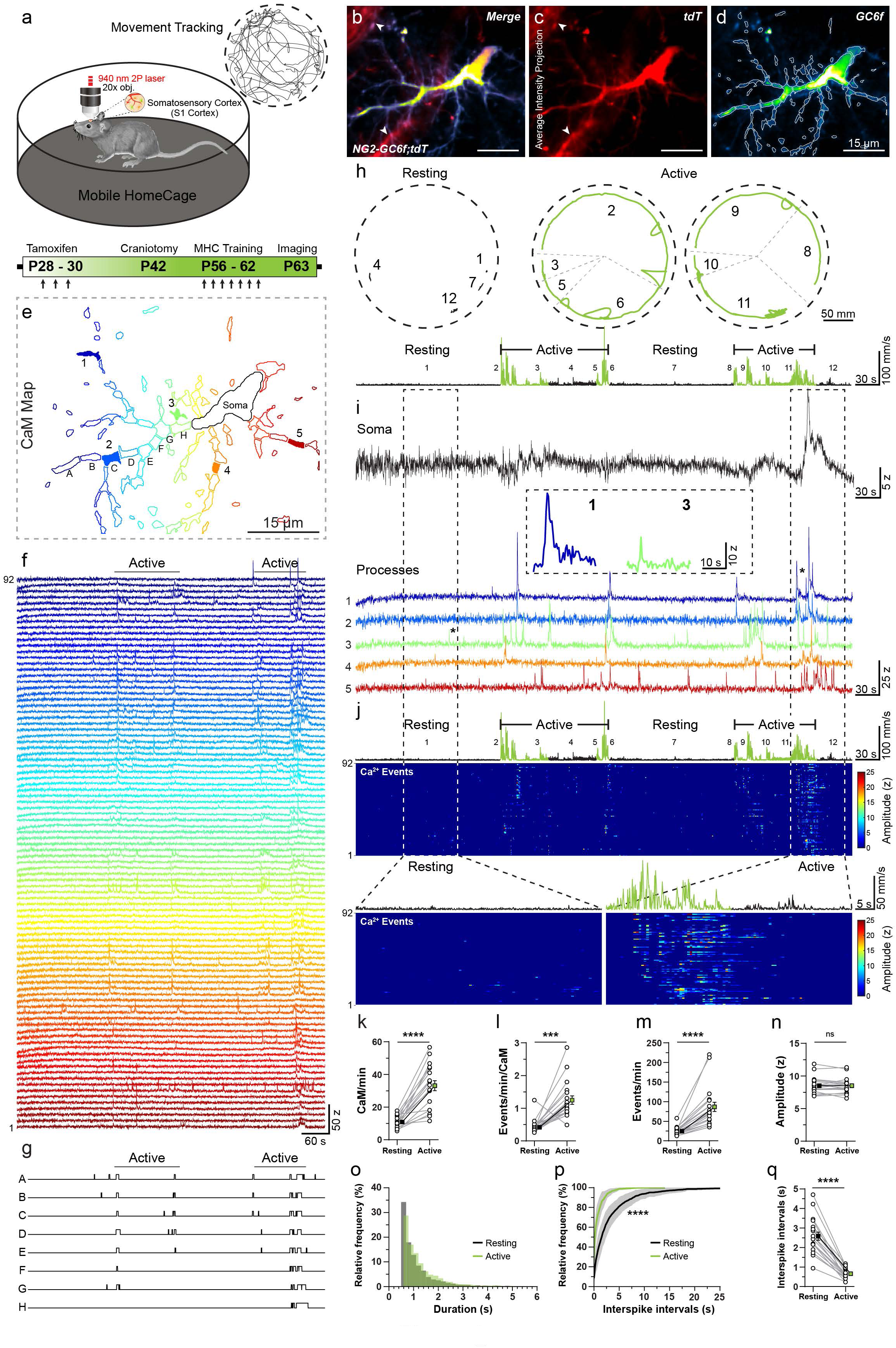
Characteristics of microdomain Ca^2+^ transients in OPCs in the cortex during baseline and activity state of mice. (**a**) Schematic representation of a head-fixed mouse exploring the Mobile HomeCage (MHC) under a 2-photon microscope for morphological and Ca^2+^ imaging of OPCs through a glass cranial window on the S1 cortex. A sample 2D tracking data representing exploratory activity of mouse in MHC is shown in the top right corner. (bottom) Experimental timeline showing induction of Cre recombination at post-natal day (P) 28, implantation of glass cranial window 2 weeks later (at P42), a one-week head-fixation training on MHC two weeks post-craniotomy (at P56), and chronic imaging of tdTomato and GCaMP6f post-training. (**b**- **d**) Pseudocolored average intensity projections of time series image stacks (x, y and t) of a recombined OPC in *NG2-GC6f;tdT* mouse co-expressing cytosolic tdTomato (tdT, **c**) and an GCaMP6f (GC6f, **d**). Map of active Ca^2+^ microdomains overlaid on GC6f image (**d**). Arrowhead indicate pericytes expressing tdT, but not GC6f (**b-c**). Scale bar, 15 µm. (**e**) Color-coded map of active Ca^2+^ microdomains (CaMs as shown in **d**) detected by the CaSCaDe algorithm over the 10-minute imaging period. Scale bar, 15 µm. (**f**) Color-matched intensity vs time Ca^2+^ traces of 92 CaMs shown in **e**. (**g**) Binary event-detection maps from neighboring CaMs labelled (A-H) in **e**. (**h**) 2D representation of the location and locomotion of the mouse in MHC arena (dashed line) during resting (black: left) or active (green: middle and right) phases. Numbers (1-12) represent distinct resting and activity bouts on activity trace (below). (**i**) Ca^2+^ transients in soma (top, black) and processes (bottom, see 1-5 color-coded in **e**) of an OPC during resting and active states of the mouse. (inset) Examples of Ca^2+^ events (asterisks on traces 1 and 3) detected in the processes. (**j**) Heat-map coded raster plots showing intensity and temporal distribution of Ca^2+^ events aligned with the associated mouse activity trace (top). Periods of exploratory locomotion activity are highlighted (green, Active). (bottom) Enlarged raster plots of intensity and temporal distribution of Ca^2+^ events during a resting phase (left, bout #1) or an active phase (right; bout #11). (**k**-**n**) Graphs comparing between the resting and active phases: number of active CaMs per minute (**k**), frequency of Ca^2+^ events per minute per CaM (**l**), frequency of Ca^2+^ events per minute (**m**), amplitude of Ca^2+^ events (**n**). (**o**) Frequency distribution histograms depicting distribution of the duration of Ca^2+^ events (in seconds) during resting (grey) and active (green) phases. (**p**) Cumulative frequency distribution plot showing distribution of interspike intervals between consecutive Ca^2+^ events during resting (black) and active (green) phases (Kolmogorov-Smirnov Test: ****p < 0.0001). (**q**) Graphs comparing between the resting and active phases average interspike intervals. All data are presented as mean ± SEM from n = 18 cells. Wilcoxon matched-pairs signed rank tests: ***p < 0.001, ****p < 0.0001; ns, not significant. mm, millimeter; s, seconds; z, z-score.

While we imaged OPC morphology and Ca^2+^ activity, we tracked the location, trajectory, and locomotion speed of the mouse in real time (Fig. 2a, h). The motion tracking data enabled us to precisely correlate microdomain Ca^2+^ activity in OPCs with the spatial location and locomotion of the mouse in MHC (Fig. 2h-j). Over a 10-minute recording interval, we observed that somatic Ca^2+^ transients in OPCs were a rare occurrence, and were exclusively seen during intense locomotion activity (Fig. 2i). Next, we found that only a small number of CaMs (10.71 ± 3.54 CaMs/min) are activated in OPCs during the ‘resting’ bouts (i.e., when mice remain at one location for more than 30s) (Fig. 2j, bouts 1, 4, 7, 12) with each CaM sparsely firing Ca^2+^ transients (0.43 ± 0.22 events/min/CaM) (Fig. 2j-l).

Conversely, when the mouse engaged in explorative bouts of locomotion (‘active’ phase), the number of activated CaMs in OPCs tripled (32.69 ± 13.58 CaMs/min) (Fig. 2j, k) and each CaM exhibited nearly 3 times more Ca^2+^ transients (1.26 ± 0.57 events/min/CaM) than during the resting phase (Fig. 2j, l and Supplementary Video 1). While the mean frequency of microdomain Ca^2+^ transients increased significantly (resting: 24.45 ± 10.69 events/min and active: 86.64 ± 52.80 events/min), there was no change in the average amplitude of Ca^2+^ events (resting: 8.52 ± 1.29 z and active: 8.49 ± 1.23 z) between the resting and the active phases (Fig. 2m, n). During locomotion however, the average duration of microdomain Ca^2+^ transient increased slightly (resting: 1.22 ± 1.49 s and active: 1.32 ± 1.43 s) (Fig. 1o). In addition, interspike interval (ISI) calculations indicate that, during the active phase, 80% of Ca^2+^ events occurred within 1.18 s of each other (Fig. 2p). On average, successive Ca^2+^ events occurred at 0.67 ± 0.27 s intervals during the active phase, but were much sparser (2.62 ± 0.92 s intervals) during the resting phase (Fig. 2q).

In neurons, excitatory post-synaptic currents (EPSCs) can be captured by GCaMP6 as fast Ca^2+^ transients (< 0.6 s)^38^. However, whether EPSCs at OPC-axon synapses generate enough Ca^2+^ flux in OPC CaMs to be visualized as ‘fast’ Ca^2+^ transients remains an open question. Using the ultrafast Ca^2+^ sensor GCaMP6f, we imaged single OPCs twice on the same day at acquisition speeds of 5 Hz (Extended Data Fig. 2a, b) and 15 Hz (Extended Data Fig. 2a, c) while mice freely engaged in explorative behavior. Indeed, we found that fast microdomain Ca^2+^ transients ranging between 330-600 ms can be detected in OPCs, when imaged at a 15 Hz sampling frequency (Extended Data Fig. 2d-g). Such fast Ca^2+^ transients (< 0.6 s) constituted about 9.7% and 7.0% of all Ca^2+^ events during resting and active phase, respectively (Extended Data Fig. 2h, i). Although the frequency of fast Ca^2+^ events (< 0.6 s) increased slightly in response to locomotion, the change in frequency of ‘slower’ Ca^2+^ events (> 0.6 s) accounted for the majority of the increase during locomotion (Extended Data Fig. 2j-l). While imaging at 15 Hz, we also saw a significant increase in the number of CaMs and the frequency of Ca^2+^ events while mice actively explored the cage, similarly to what was observed at 5 Hz. Taken together, these results suggest that *in vivo* OPCs exhibit a wide variety of fast and slow Ca^2+^ signals, which are mostly restricted to CaMs in the processes, and that OPC Ca^2+^ transients increase significantly during active explorative behavior.

### Microdomain Ca^2+^ signals correlate with the fate of OPCs

Ca^2+^ can regulate all aspects of the OPC fate such as proliferation, differentiation and programed cell death. As OPCs divide into daughter cells or differentiate into OLs, they might undergo profound changes in the cellular connectivity, neurotransmitter receptor expression, and their overall response to neuronal activity. To study whether OPCs generate distinct Ca^2+^ signals as they proliferate and differentiate, we used *NG2-GC6f;tdT* mice to concurrently track the lineage progression (through changes in the cell morphology) of OPCs and their Ca^2+^ activity at each stage of OPC fate. We imaged tdTomato and GCaMP6f signals in individual OPCs in the S1 cortex every 2-4 days over a period of 2 weeks (Fig. 3a- f). A detailed analysis of Ca^2+^ signals from a typical cell (out of 35 tracked cells) showed that OPCs maintained a very stable microdomain Ca^2+^ activity (Fig. 3g, h) with slight morphological change over several days (Day 0, 2, 6, and 9; Fig. 3a-d). During this stable period, OPCs remained responsive to locomotion-induced increases in the ongoing neuronal activity, and increased both the number of active CaMs (Day 0, 2, 6, and 9; Fig. 3g) and frequency of events (Fig. 3h) approximately 3-fold. As the OPC began to differentiate into an OL, we noted prominent morphological changes as seen by a circular soma and increased cellular complexity (see Fig. 1d, e; Fig. 3e, f: Day 12 and 15). Once the OPC transitioned into a differentiated state (Days 12, 15), there were no significant changes in the number of CaMs (Fig. 3g) or the frequency of Ca^2+^ events (Fig, 3h) in response to locomotion. Also, ISIs between the resting and active phases of exploratory behavior indicate a loss of responsiveness to the locomotion once the OPC (Days 0, 2, 6, 9: Fig. 3i) differentiated into an OL (Days 12 and 15: Fig. 3i).

**Figure 3.**
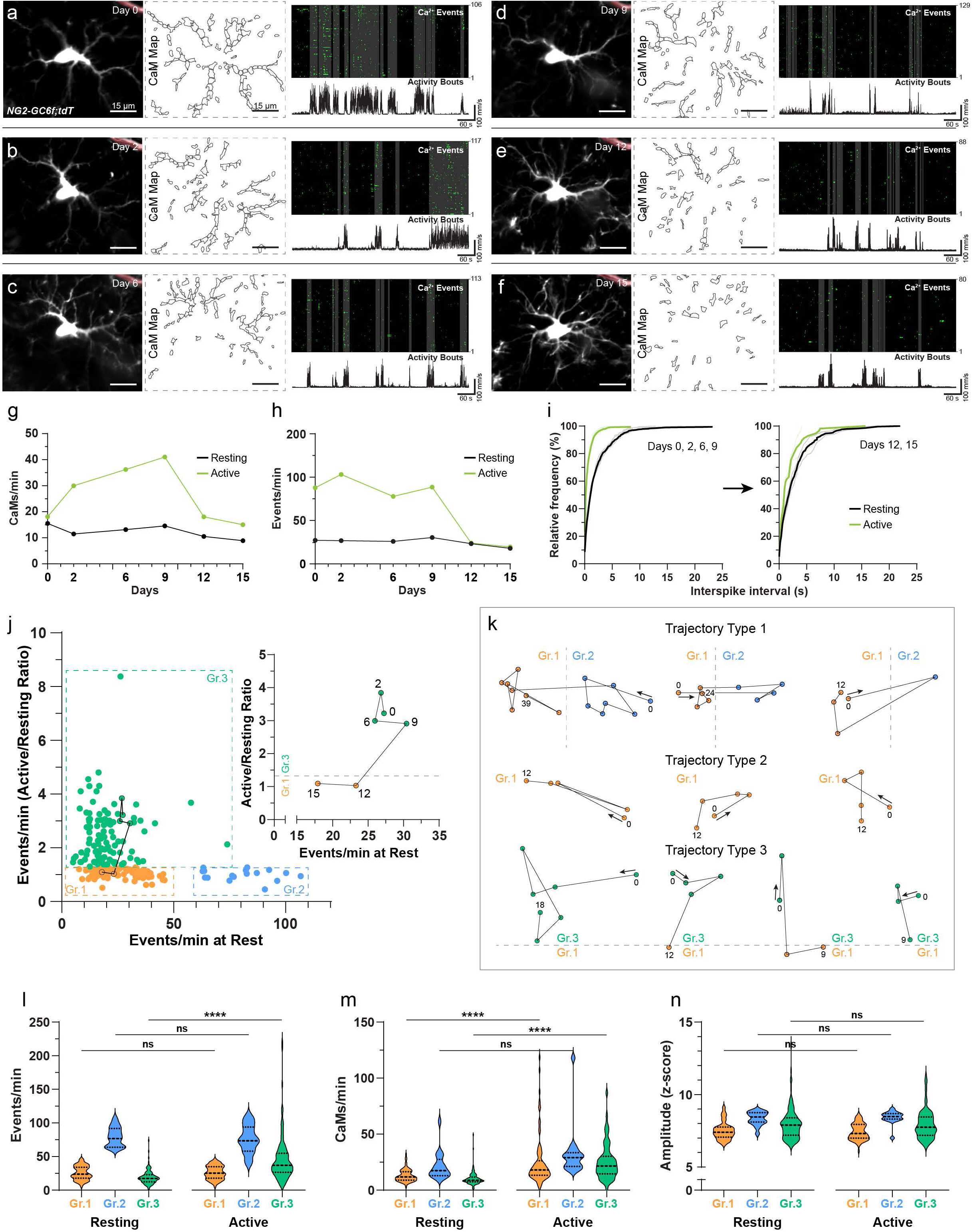
Evolution of Ca^2+^ signals as OPCs progress in their fate. **(a)** (left) Average intensity time-series projection of tdTomato reporter expressed by an OPC in the S1 cortex of *NG2-GC6f;tdT* mouse at the start (Day 0) of the chronic imaging session, and (middle) its associated microdomain Ca^2+^ activity (CaM) map. (right) Binarized raster plot showing detected Ca^2+^ events in all 106 CaMs (top) and locomotion activity trace of the mouse during a 10-min imaging period (bottom). Vertical grey bars in the raster plot represent locomotion bouts. A recombined pericyte next to a blood vessel was used to precisely relocate and align the same cell over 2 weeks imaging period (top right, marked as light red). (**b**-**f)** Morphological changes, CaM maps, microdomain Ca^2+^ transients and mouse locomotion activity bouts of the same OPC shown in **a** over a period of 2 weeks at (**b**) Day 2, (**c**) Day 6, (**d**) Day 9, (**e**) Day 12, (**f**) and Day 15. (**g**, **h**) Graphs showing changes in the number of CaMs (**g**), and the frequency of Ca^2+^ events (**h**) during resting and active state of the mouse as the OPC progress towards differentiation into oligodendrocyte (shown in **a-f**; Day 0 – Day 15). (**i**) Cumulative frequency distributions of interspike intervals during the resting (black) and active (green) phases. Grey lines correspond to individual time points, and thicker black (resting) and green (active) lines represent average frequency distributions of all cells. Note the shift in interspike intervals between the first 4 (left; Day 0 – Day 9) and the last 2 (right; Day 12, and day 15) imaging sessions. (**j**) Graphs showing the ratio between the frequency of Ca^2+^ events during resting and active phases in each cell as a function of its “baseline” activity (i.e., number of events per minute at rest). Dashed boxes delimit 3 distinct groups (Gr.1-3) of cells. (right) a constellation plot highlights the Ca^2+^ activity state at every time point (Day 0 - Day 15) as the OPC (shown in **a**-**f**) undergoes fate progression. In total 214 time-points were imaged from 9 mice. (**k**) Constellation plots illustrates 3 types of trajectories (type 1-3) that cells took in the 2D space over time. Individual dots in the constellation corresponds to unique imaging time point of the same cell, and the numbers indicate endpoint of imaging. Arrows indicate the direction of the trajectory and the first imaging point of the cell. Dashed lines show the transitioning of cells form one group to the other. (**l**-**n**) Violin plots comparing between resting (left) and active (right) phases, the frequency of Ca^2+^ events (**l**), the number of active CaMs (**m**), and the amplitude of Ca^2+^ events (**n**) when cells are in group 1 (orange, n = 88), group 2 (blue, n = 17) and group 3 (green, n = 109). Kruskal-Wallis test performed with Dunn’s multiple comparisons: ****P<0.0001; ns, not significant.

Once OPCs undergo Cre-mediated recombination in the *NG2-GC6f;tdT* mouse line, tdTomato and GCaMP6f are permanently expressed even after OPCs divide or differentiate. Therefore, it is likely that the OPCs we imaged were scattered across the distinct stages of the OL lineage. To study the dynamics of the fate of OPCs, we clustered all 214 cells we imaged based on the frequency of Ca^2+^ events when mice were in resting state (x-axis), and the ratio of frequency of Ca^2+^ events during resting and active state (A/R ratio; y-axis). When the Ca^2+^ events frequency increased by >30% in response to locomotion i.e. A/R ratio of 1.3, cells were considered as responsive to locomotion (Fig. 3j). Based on this criterion, we intuitively classified OLCs into three broad groups (Fig. 3j-n): (**1**) cells with low number of baseline Ca^2+^ events (< 50 events/min) that are unresponsive to locomotion (n = 88 cells; mean A/R ratio = 1.04 ± 0.16) (Fig. 3j, l, m); (**2**) cells with high frequency of baseline Ca^2+^ transients (> 50 events/min), but don’t further enhance their Ca^2+^ activity in response to locomotion (n = 17 cells; mean A/R ratio = 0.99 ± 0.21) (Fig. 3, l-n); and (**3**) cells with highly variable baseline Ca^2+^ transients that respond robustly to locomotion (n = 109 cells; mean A/R ratio = 2.43 ± 1.02) (Fig. 3j, l, m). A *constellation plot* shows the evolution of Ca^2+^ activity during the resting and locomotion phase as an OPC undergoes lineage progression (Fig. 3j, inset: Days 0, 2, 6, 9, 12, and 15). After mapping constellation plots of OLCs tracked for several days (Fig. 3k), we found that OPCs in Group 1 and Group 2 stayed in their respective group for several days, and could migrate back and forth between both the groups (Fig. 3k, Trajectory Type 1). We noted that OPCs cell division occurred mainly in Group 1 (5 division events out of 88 cell recordings) and that dividing cells toned down their Ca^2+^ activity (Extended Data Fig. 3a, b). When OPCs split into two daughter cells, the total number of CaMs and the frequency of Ca^2+^ events of both the daughter cells together were equivalent to that of the original cell (Extended Data Fig. 3c-e), thereby implying that daughter cells can inherit cellular connectivity and the repertoire of neurotransmitter receptors from their mother cell (Supplementary Video 2). In addition, the mother and daughter cells often don’t respond to locomotion (Extended Data Fig. 3f). In general, Group 1 cells with OL-like morphology have a stable number of CaMs and Ca^2+^ events, and stay within the group (Fig. 3k, Trajectory Type 2; Extended Data Fig. 3g, h). Although these cells show slight increase in the number of CaMs (resting: 12.62 ± 5.66 CaMs/min and active: 16.18 ± 3.92 CaMs/min), frequency of Ca^2+^ events are low and cells don’t show increase in Ca^2+^ events in response to locomotion (resting: 25.99 ± 10.13 and active: 26.98 ± 9.53 events/min) (Extended Data Fig. 3j-l, and Supplementary Video 3). In contrast, OLCs in Group 3 are in their most dynamic state, stay within Group 3 for several days or weeks, then migrate towards Group 1 and stop responding to locomotion (Figure 3k, Trajectory Type 3). Over time, Group 3 cells differentiated into OLs, migrated (out of imaging frame) or died. In conclusion, OPCs exhibit unique Ca^2+^ signatures at different stages during their lineage progression. They lose the capability to integrate locomotion-evoked neuronal activity prior to division, or are highly responsive to it before differentiating into OLs.

### Noradrenaline release induces microdomain Ca^2+^ transients in OPCs

In rodents, locomotion activates neurons in the locus coeruleus (LC), which in turn release norepinephrine (NE) across several brain regions, including the cortex^39^. Since axonal projections of LC neurons in the cortex are very thin and profusely branched, cytosolic variants of GCaMPs don’t reach high enough concentration in axons for a reliable measurement of Ca^2+^ signals in *vivo*^40^. We therefore used a membrane anchored, sensitive variant of GCaMP6 (mGCaMP6s) to image Ca^2+^ signals in LC axons. We cross-bred *Dbh- Cre* mice^41^ with our newly generated Cre-dependent mGCaMP6s mouse line to generate double transgenic mice (*Dbh-Cre;Rosa26-LSL-mGCaMP6s***; *Dbh-mGC6s***) (Fig. 4a; Extended Data Fig.1a). The immunohistochemical analysis in the S1 cortex of adult *Dbh- mGC6s* mice confirmed that all axons expressing mGCaMP6s (labeled with anti-eGFP antibody) were also positive for tyrosine hydroxylase, an enzyme involved in NE synthesis and labelling adrenergic neurons in the cortex (Fig. 4b). Similar to the *in vivo* Ca^2+^ imaging experiments in OPCs, we implanted a chronic cranial window on the S1 cortex of 6-8 weeks old *Dbh-mGC6s* mice (time line: Fig. 2a). To assess whether locomotion bouts during explorative behavior in the MHC can engage LC neurons, we imaged Ca^2+^ transients of LC axonal projections in the S1 cortex (Fig. 4c, d). We found that when the mouse was in a ‘resting’ state, LC neurons fired occasionally (16.72 ± 13.4 events/min) (Fig. 4e-g). However, as soon as mice started to actively explore, as seen by locomotion bouts (Fig. 4f), the firing rate of LC neurons increased by 3.5 times (57.91 ± 36.9 events/min), but there was no change in the amplitude of Ca^2+^ events between the resting and active phases (Figure 4g, h; Supplementary Video 4). Additionally, cortical projections of LC neurons mostly fired in sync with the locomotion bouts (Fig. 4e, f), and the maximal response was reached within 300 ms from the initiation of locomotion (Fig. 4i). Notably, the latency of the maximal response (i.e. maximum number of active CaMs) in OPC was about 3.6 s longer than the neuronal response (Fig. 4i; also see Fig. 2j – zoom-in of active bout).

**Figure 4.**
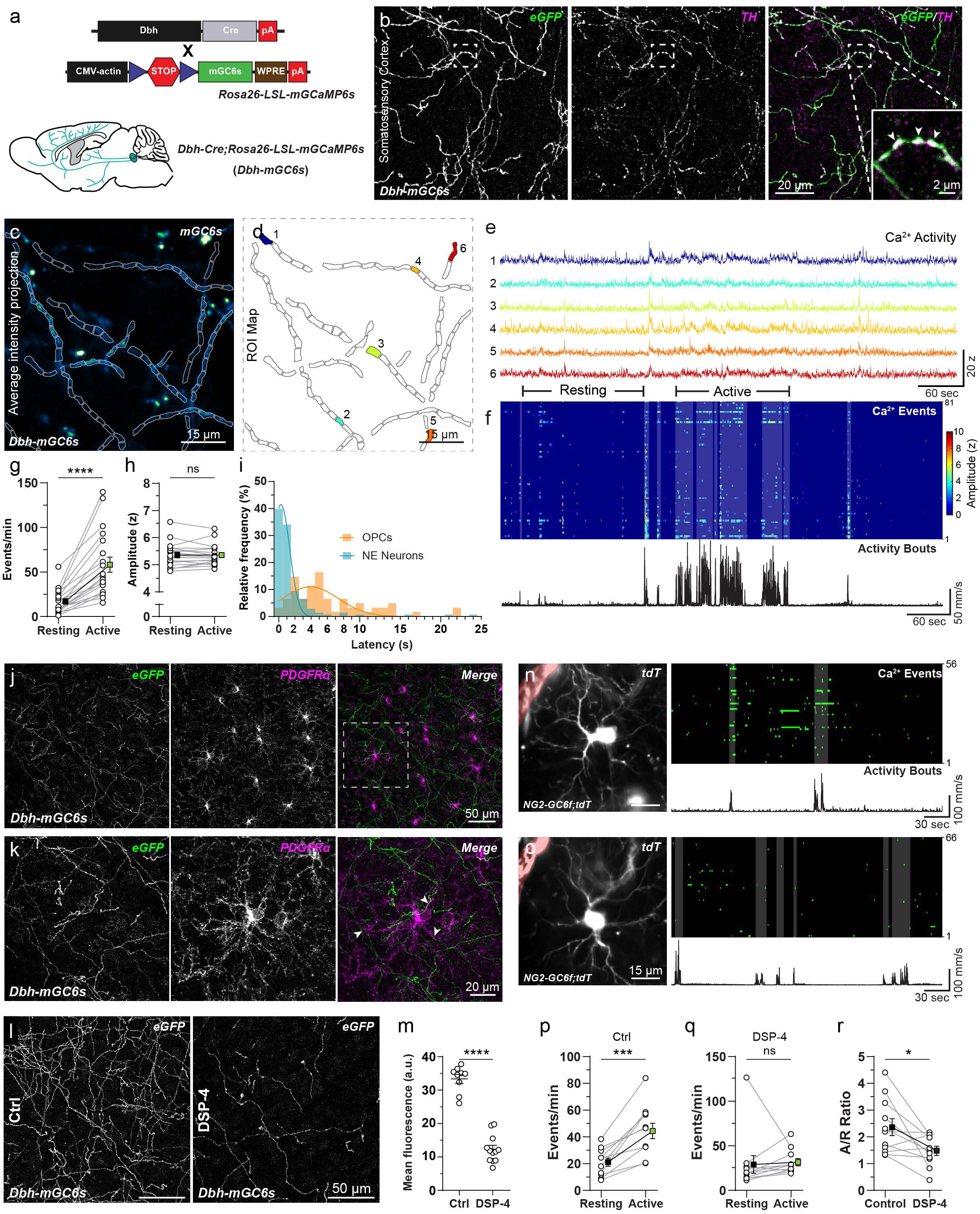
Locomotion-induced Ca^2+^ transients in OPCs is mediated by norepinephrine. **(a)** (top) Cartoon showing a transgenic strategy to express membrane anchored variant of GCaMP6s (mGCaMP6s) in noradrenergic (NA) sneurons in *Dbh-Cre;Rosa26-LSL- mGCaMP6s* (*Dbh-mGC6s*) mice. (bottom) mouse brain cartoon showing the distribution of noradrenergic projections from the locus coeruleus (LC) to the cortex. (**b**) Maximum intensity projection image of a confocal z-stack confirming expression of mGCaMP6s (green) by NA axonal projections (magenta) in the S1 cortex of *Dbh-mGC6s* mice as seen by immunohistochemical analysis using antibodies against eGFP and tyrosine hydroxylase (TH) respectively. (inset) Arrowheads in the high-magnification image of the boxed area highlights norepinephrine (NE) release site (varicosities) on the NA axons. Scale bars, 20 µm and 2 µm (inset). (**c, d**) Average intensity time-series projection (pseudocolored) of NA fibers in the S1 cortex of *Dbh-mGC6s* mice, overlayed with active region with Ca^2+^ signals (**c**), and color-coded map of detected Ca^2+^ regions of interest (ROIs) (**d**), scale bar, 15 µm. (**e**) Example traces showing Ca^2+^ signals of the 6 ROIs shown in **d**. (**f**) Heat map raster plot of the intensity and temporal distribution of Ca^2+^ events (top), aligned with the associated mouse locomotion activity (black trace, bottom). Grey boxes in raster plot show correlation between locomotion activity and Ca^2+^ signals. (**g**, **h**) Graphs comparing frequency (**g**) and amplitude (**h**) of Ca^2+^ events in NA fibers during resting and active phases (n = 19; Wilcoxon matched-pairs signed rank tests: ****P < 0.0001; ns, not significant). (**i**) Frequency distribution histograms of the time-lag between the initiation of locomotion and Ca^2+^ activity in NA neurons (cyan) and OPCs (orange). (**j)** Maximum intensity projection image of a confocal z-stack of the cortex from *Dbh-mGC6s* mice stained for eGFP (green) and PDGFRα (magenta). Scale bar, 50 µm. (**k**) High-magnification image of PDGFRα+ OPC and NA fibers in the boxed area in **j**. Arrowheads highlight areas of close contact between PDGFRα+ OPC and eGFP+ NA fibers. Scale bar, 20 µm. (**I)** Maximum intensity projection image of a confocal z-stack of control (Ctrl) and DSP-4 treated *Dbh-mGC6* mice stained for eGFP (green). Scale bar, 50 µm. (**m**) Graph showing the mean fluorescence intensity of eGFP in coronal sections of control (n = 9 sections) and DSP-4 treated (n = 12 sections) *Dbh-mGC6s* mice (Unpaired t-test: ****P < 0.0001). (**n, o**) (left) Average intensity projection of time-series of an OPC from a *NG2-GC6f;tdT* mouse imaged prior (**n**) and after (**o**) DSP-4 treatment. (right) Raster plots showing Ca^2+^ events (top) and locomotion activity (bottom). A pericyte expressing tdT (light red) on the top left corner was used to precisely locate and align OPCs across imaging sessions (**n, o**). Scale bar, 15 µm. (**p**, **q**) Graphs showing frequency of Ca^2+^ events during the resting & active phases before (**p**) and after (**q**) treatment with DSP-4. n = 11 cells; Paired t-test: ***P = 0.0004 (control) and Wilcoxon matched-pairs signed rank tests: P = 0.1016 (DSP-4). (**r**) Graph showing changes in the ratio between the number of Ca^2+^ events per minute detected during resting and active phases (A/R ratio) before and after treatment with DSP-4 (n = 11 cells; Paired t-test: *P = 0.0187). For all graphs data are presented as mean ± SEM. Ctrl, control.

To understand the anatomical interaction between OPCs and the cortical projections of LC neurons, we immunohistochemically visualized LC fibers expressing mGCaMP6s and OPCs in the S1 cortex of *Dbh-mGC6s* mice using antibodies against eGFP (mGCaMP6s is a modified eGFP) and PDGFRα, respectively (Fig. 4j, k). These experiments revealed a close contact between the LC fibers and OPC processes, indicating that OPCs are well placed to sense and respond to NE released by the LC fibers. To further ensure that activation of LC and concomitant NE release in response to locomotion indeed induces Ca^2+^ transients in OPCs, we specifically depleted the LC projections in the cortex of *Dbh-mGC6s* mice by injecting a single dose of the neurotoxin DSP4 (50mg/kg)^42^. 3-4 days after DSP4 injection, we performed histological analysis to visualize LC fibers expressing mGCaMP6s, and found that almost two thirds (63.12%) of the LC projections in the cortex were successfully ablated (Fig. 4l, m). Next, we imaged Ca^2+^ transients in single OPCs in the S1 cortex of *NG2- GC6f;tdT* mice (Fig. 4n), then injected the mice with DSP4 (50mg/kg). 3-4 days after the DSP-4 injection, we imaged Ca^2+^ transients in the same set of OPCs (Fig 4o). As expected, the frequency of Ca^2+^ events in these OPCs originally increased in response to locomotion, but this effect was abolished in the same set of OPCs after the ablation of the LC fibers (Fig 4p, q), and the A/R ratio dropped significantly (Fig. 4r). Hence, we conclude that locomotion induced Ca^2+^ transients in OPCs are dependent on the activation of LC neurons and NE release.

### Activation of alpha-adrenergic receptors on OPCs increases Ca^2+^ transients

OPCs express a wide variety of ionotropic and metabotropic receptors for neurotransmitters and neuromodulators including glutamate, GABA, ATP, ACh and NE. To pharmacologically dissect the source of Ca^2+^ transients in OPC, we performed 2-photon Ca^2+^ imaging in the S1 cortex in acute brain slices derived from 6-12 weeks old *NG2-mG6s;tdT* mice (Fig 1. F, g and Extended Data Fig. 1). Like *in vivo*, baseline Ca^2+^ transients of OPCs in acute brain slices were spatially restricted to discrete microdomains (Fig. 5a, b, and Extended Data Fig. 4a-d). However, the number of activated CaMs (1.89 ± 1.78 CaMs/min) and the frequency of Ca^2+^ transients at baseline in OPCs (3.79 ± 3.8 events/min) were very low compared to the baseline activity seen *in vivo* (see Fig. 2k-m). These baseline Ca^2+^ transients persisted in OPCs when the neuronal activity was blocked by a voltage-gated sodium channel blocker tetrodotoxin (0.5 µM; TTX) (Extended Data Fig. 4a-e). There were almost no changes in the number of CaMs (TTX: 1.92 ± 1.46 CaM/min) or the frequency of Ca^2+^ events (TTX: 3.64 ± 3.65 events/min), but a slight increase in the amplitude of Ca^2+^ events in the presence of TTX (Supplementary Fig. 4e-h). In addition, only a small fraction of the baseline Ca^2+^ transients in OPCs were blocked by ionotropic and metabotropic glutamate receptors (Glut- B: TTX, CNQX, D-AP5, MCPG) (Extended Data Fig. 4i-l) or GABA receptor (Gaba-B: TTX, Gabazine, CGP55845) antagonists (Extended Data Fig. 4m-p), suggesting that baseline Ca^2+^ transients in OPCs are not dependent on the spontaneous release of glutamate and GABA at OPC-axon synapses. Thus, we suggest that OPCs can generate low frequency spontaneous microdomain Ca^2+^ events, which are driven by cell-intrinsic mechanisms possibly similar to those previously described for cortical astrocytes^37^.

**Figure 5.**
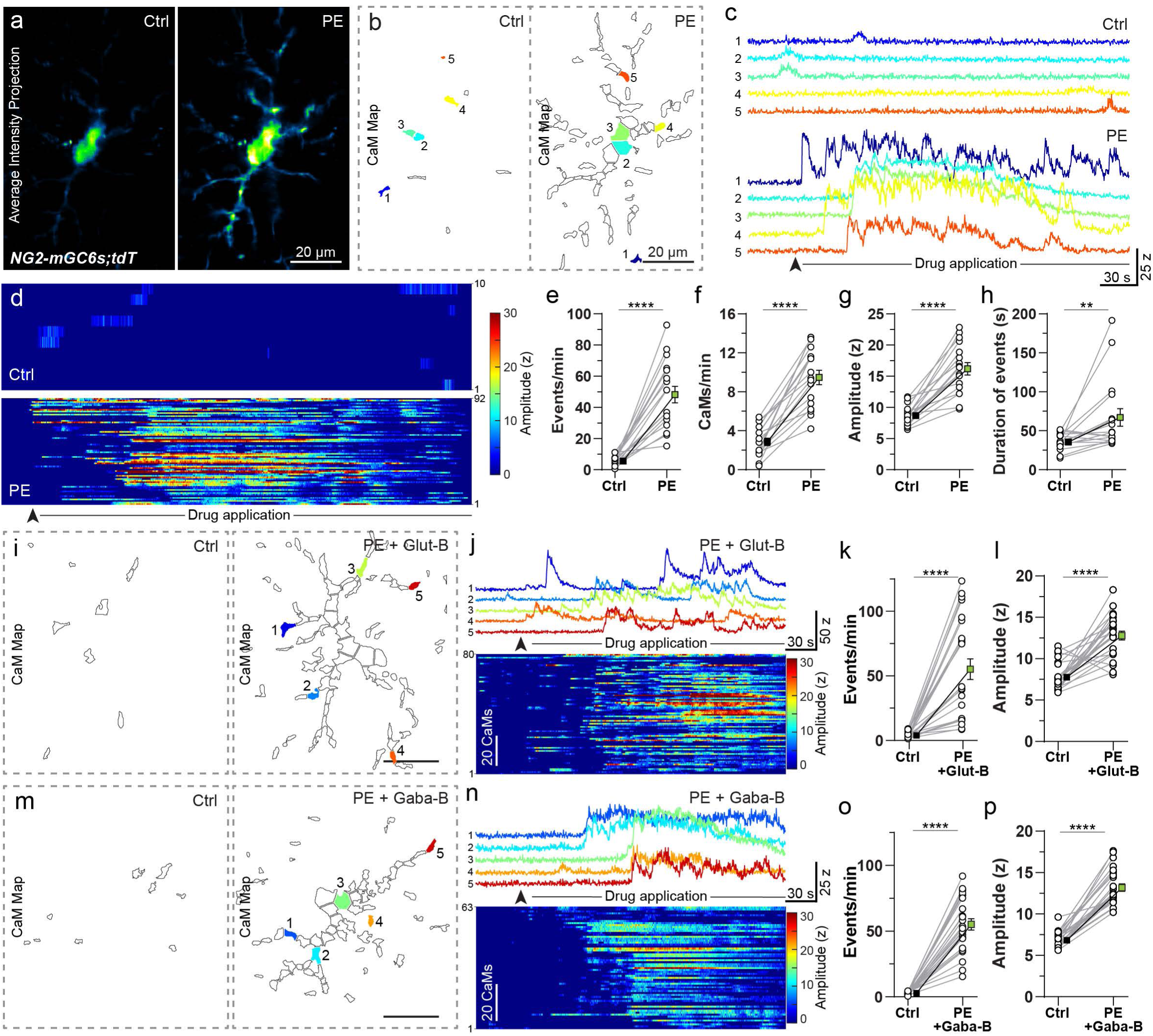
Direct activation of α1 adrenergic receptors on OPCs induces Ca^2+^ transients. (**a**) Pseudocolored average intensity time-series projection of mGCaMP6s signal in an acute brain slice OPC from a *NG2-mGC6s;tdT* mouse in control (Ctrl) conditions (TTX, 0.5 µM; left) and after bath application of phenylephrine (PE; 10 μM + TTX; 0.5 µM). (**b**) CaM map recorded in the Ctrl (left) and after PE application (right) (**c**) Intensity vs time Ca^2+^ traces from 5 CaMs in Ctrl (top) and after PE application (bottom) corresponding to colors in **b**. (**d)** Heat-map raster plots of the intensity and temporal distribution of Ca^2+^ events in the Ctrl (top) and after PE application (bottom). (**e**-**h)** Graphs showing changes in frequency of Ca^2+^ events (n = 17 cells; Paired t-test: ****P<0.0001) (**e**), number of active CaMs per minute (n = 17 cells; Paired t-test: ****P<0.0001) (**f**), average amplitude (z-scores) of Ca^2+^ events (n = 17 cells; Paired t-test: ****P<0.0001) (**g**), and mean duration of Ca^2+^ events (n = 17 cells; Wilcoxon matched-pairs signed rank test: **P<0.0011) (**h**) in Ctrl and after PE application. (**i**) Map of CaMs in control (Ctrl – Glut-B: TTX, 0.5 µM; CNQX, 10 µM; AP5,50 µM; MCPG, 10 µM; left) and after PE + Glut-B application. (**j**) Intensity vs time Ca^2+^ traces from 5 CaMs after application of PE + Glut-B corresponding to colors in **i** (top). Heat-map raster plots of the intensity and temporal distribution of Ca^2+^ events after PE + Glut-B application (bottom). (**k**, **l**) Graphs showing changes in frequency (n = 24 cells; Paired t-test: ****P<0.0001) and average amplitude (n = 24 cells; Paired t-test: ****P<0.0001) of Ca^2+^ events in Ctrl and after application of PE + Glut-B. (**m**) Map of CaMs in control (Ctrl – Gaba-B: TTX, 0.5 µM; CGP 55845, 5 µM; Gabazine, 5 µM; left) and after PE + Gaba-B application. (**n**) Intensity vs time Ca^2+^ traces from 5 CaMs after PE + Gaba-B application corresponding to colors in **m** (top). Heat-map raster plots of the intensity and temporal distribution of Ca^2+^ events after PE + Gaba-B application (bottom). (**o**, **p**) Graphs showing changes in frequency (n = 24 cells; Paired t-test: ****P<0.0001) and amplitude (n = 24 cells; Wilcoxon matched-pairs signed rank test: ****P<0.0001) of Ca^2+^ events in Ctrl and after application of PE + Gaba-B. All data are presented as mean ± SEM. Scale bars, 20 µm.

To determine if OPCs express functional adrenergic receptors, and whether direct activation of these receptors can induce microdomain Ca^2+^ transients, we bath-applied a potent alpha1 adrenergic receptor (α1-AR) agonist, phenylephrine (10µM; PE) in the presence of neuronal action potential blocker TTX (0.5 µM) (Fig 5a, b; Supplementary Video 5). We observed that activation of α1-AR on OPCs induced a large increase in the number of CaMs (3.3x), frequency (8.7x), amplitude (1.9x) and duration (1.9x) of Ca^2+^ events when compared to the baseline activity (in TTX, 0.5 µM) (Fig. 5e-g). To further validate that the increase in Ca^2+^ transients is due to the direct activation of α1-AR on OPCs (and not through an indirect activation of neurons, leading to the release of glutamate or GABA), we bath-applied PE in the presence of a cocktail of drugs composed of a neuronal activity blocker (TTX) combined with either **1)** ionotropic and metabotropic glutamate receptors antagonists (Glut-B: CNQX, D-AP5, MCPG) (Fig. 5i-k) or **2)** GABA (Gaba-B: Gabazine, CGP55845) receptors (Fig. 5m- p) antagonists. In the presence of these blockers, PE still evoked a large increase in the frequency and amplitude of Ca^2+^ events in OPCs (Fig. 5k, l, o, p). These experiments confirm that OPCs indeed express α1-ARs, and that a direct activation of these receptors can evoke microdomain Ca^2+^ transients in OPCs.

### All three sub-type of alpha-adrenergic receptors are expressed on OPCs

There are 3 subtypes of α1-ARs, namely α1a, α1b and α1d, widely expressed in different brain areas^43^. All α1-ARs are coupled to G_q_-type GPCRs and upon activation lead to cytosolic Ca^2+^ increase through the opening of IP_3_Rs on the endoplasmic reticulum (ER). To identify which α1-ARs subtypes are expressed by the OLCs, we performed simultaneous immunohistochemistry and single-RNA-molecule fluorescent in-situ hybridization (sm-FISH) in the S1 cortex of adult wildtype mice (Fig 6a-c). We labeled OLCs using either an in-situ probe or an antibody for pan oligodendrocyte lineage marker Olig2. All three α1-AR subtypes were labeled with their respective in-situ probes, and nuclei of all cells were labeled with DAPI (Fig 6a). Our quantitative analysis revealed that all three sub-types of α1- ARs are expressed in OLCs, but at variable levels (Fig. 6d-e). Cells with >1 fluorescent puncta for a given α1-AR subtype in the close vicinity of their nucleus were considered positive for that α1-AR subtype (see Methods). Based on this criterion, about 30% and 64% of cortical Olig2+ OLCs expressed Adra1a and Adra1b α1-AR subtypes respectively (Fig. 6d, e). Due to incompatibility of Olig2 and Adra1d in-situ probes, we instead used an Olig2 antibody to label OLCs, and found that about 45% of cortical OLCs expressed Adra1d (Fig 6e). Next, to examine whether the activation of α1a, α1b and α1d AR subtypes can enhance intracellular Ca^2+^ in OPCs, we performed 2-photon Ca^2+^ imaging in the S1 cortex of acute brain slices from 6-12 weeks old *NG2-mG6s;tdT* mice. Due to a lack of sub-type specific α1- AR agonists, we used sub-type specific α1-ARs antagonists, and studied the level of suppression of PE-evoked Ca^2+^ transients in OPCs in the presence of these antagonists. We found that the baseline Ca^2+^ transients in OPCs persisted in the presence of TTX and the sub-type specific α1a (RS17035; 40 μM), α1b (Chloroethylclonidine, CEC; 30 μM) and α1d (BMY7378; 40 μM) antagonists (Fig. 7c, g, k), and suggest that activation of α1-ARs doesn’t contribute to the generation of spontaneous Ca^2+^ in OPCs. When we bath-applied PE in the presence of α1a, α1b and α1d antagonists, OPC showed only 102%, 95%, and 33% increase in the frequency of microdomain Ca^2+^ events respectively (Fig. 7e, i, m), with a slight increase in the amplitude of Ca^2+^ events for α1b (23%), but not for α1a and α1d α1- ARs (Fig. 7f, j, n). This is in contrast to the 866% increase in the frequency and 186% increase in amplitude of Ca^2+^ events observed in OPCs when PE (+TTX) was bath-applied to the slices without α1-ARs antagonists (Fig. 5 e, f). Taken together, these results further strengthen our conclusion that OPCs express all three subtype α1-ARs and that activation to these receptors can induce intracellular Ca^2+^ increase in CaMs.

**Figure 6.**
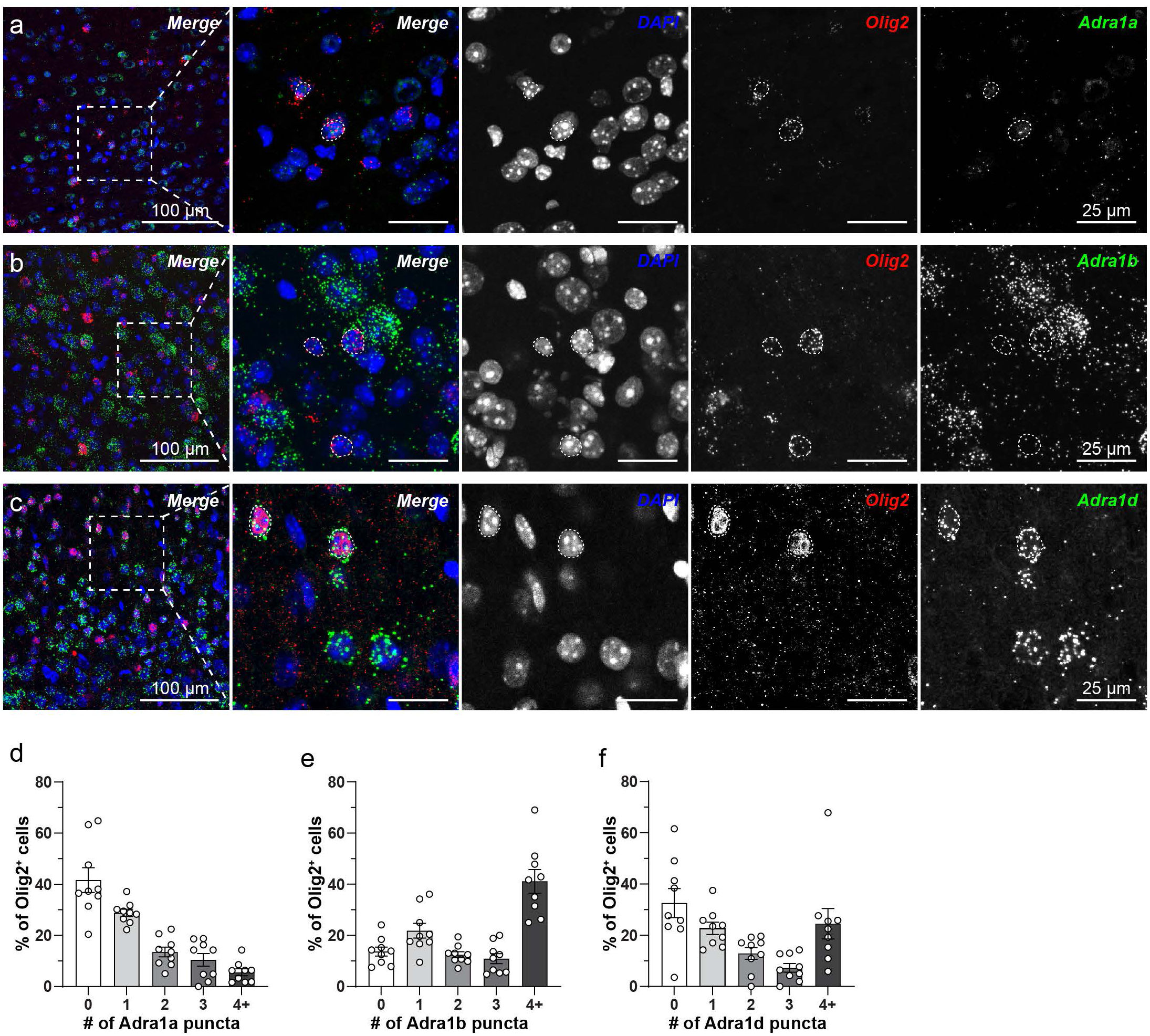
Oligodendrocyte lineage cells express all three subtypes of α1 adrenergic receptors. (**a**) (left) low and high-magnification (boxed area) maximum intensity projected confocal z- stack images of cortical brain sections labeled with a pan-nuclear maker DAPI (blue), a sm- FISH probe for Olig2 to mark all oligodendrocyte lineage cells (red), and a sm-FISH probe to detect Adra1a (green). (**b**) Maximum intensity projected z-stack images of cortical brain sections labelled with DAPI (blue), sm-FISH probe for Olig2 (red), and sm-FISH probe for Adra1b (green). (**c**) Maximum intensity projected z-stack images of cortical brain sections labelled with DAPI (blue), anti-Olig2 antibody (red), and sm-FISH probe for Adra1d (green). In all images, dotted circular outlines highlight nuclei of Olig2^+^ cells. (**d**-**f**) Relative frequency distribution of the number of Adra1a+ (**d**), Adra1b+ (**e**) and Adra1d+ (**f**) puncta in oligodendrocyte lineage cells (n = 9 slices from 3 mice). Cells with >1 puncta were considered positive for corresponding α1 adrenergic receptor sub-type. Data are presented as mean ± SEM. Scale bars, 100 µm (left) and 25 µm (boxed area).

**Figure 7.**
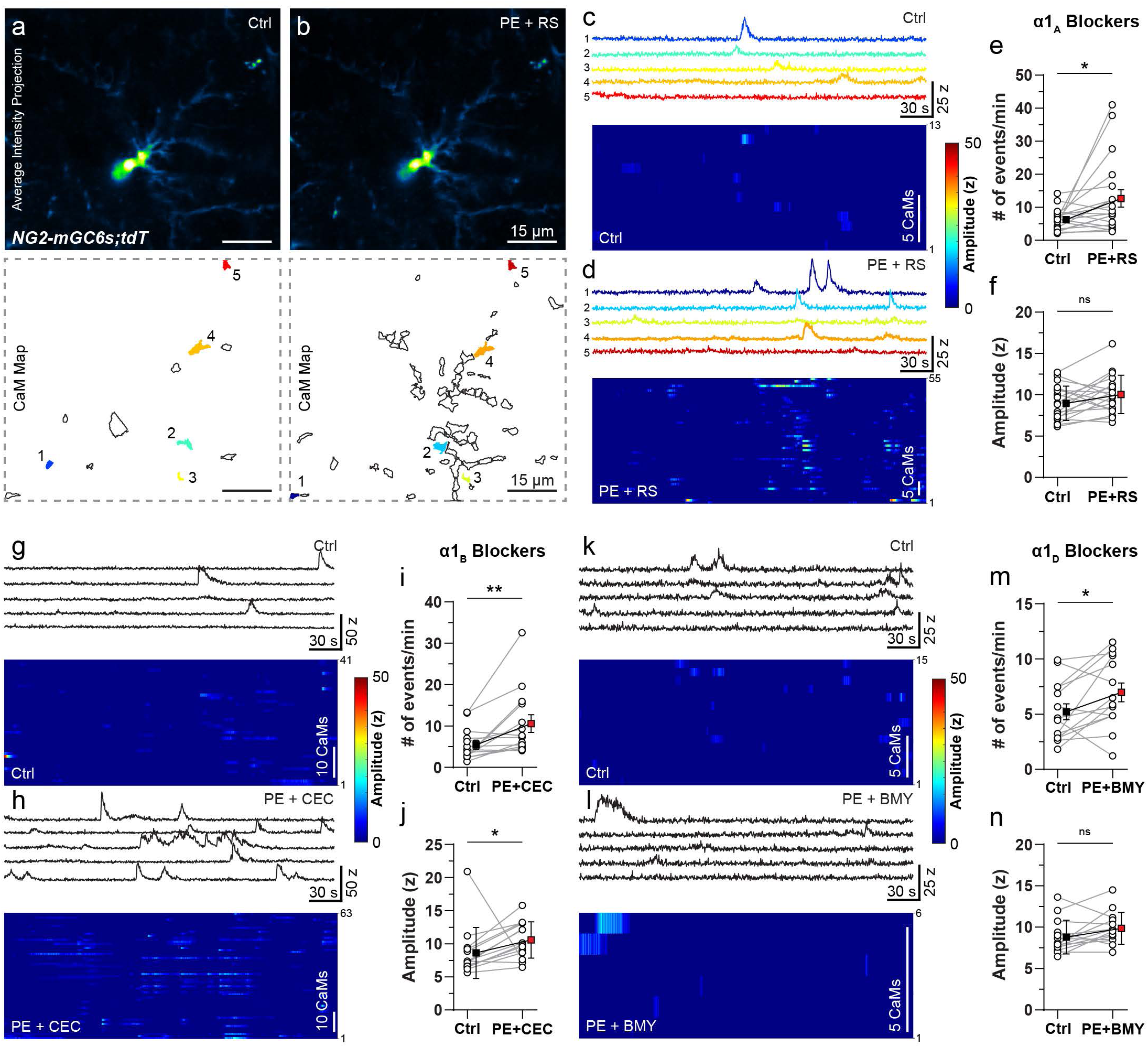
Activation of all three subtypes of α1 adrenergic receptors on OPCs induces Ca^2+^ transients. **(a, b)** Average intensity time-series projection (pseudocolored) of mGCaMP6s in an acute brain slice OPC from *NG2-mGC6s;tdT* mouse in control (Ctrl – TTX, 0.5 µM and α1a antagonist RS17053, 40 µM) (**a**, top) and after bath application of phenylephrine with RS17053 (PE + RS; 10 μM + 40 µM) (**b**, top). CaM map recorded in Ctrl conditions (**a**, bottom) and after PE + RS application (**b**, bottom). (**c, d**) Intensity vs time Ca^2+^ traces from 5 CaMs in Ctrl (**c**, top) and after PE + RS application (**d**, top) corresponding to colors in **a** and **b** respectively. Heat-map raster plots of the intensity and temporal distribution of Ca^2+^ events in the Ctrl (**c**, bottom) and after PE + RS application (**d**, bottom). (**e, f**) Graphs showing changes in frequency (n = 19 cells; Wilcoxon matched-pairs signed rank test: P=0.0181) (**e**), and average amplitude (n = 19 cells; Paired t-test: P=0.0627) of Ca^2+^ events (**f**). (**g, h**) Intensity vs time Ca^2+^ traces from 5 CaMs in control (Ctrl – TTX, 0.5 µM and α1b antagonist chloroethyclonidine (CEC), 30 µM) (**g**, top) and after PE + CEC application (**h**, top). Heat- map raster plots of the intensity and temporal distribution of Ca^2+^ events in the Ctrl (**g**, bottom) and after PE + CEC application (**h**, bottom). (**i, j**) Graphs showing changes in frequency (n = 14 cells; Wilcoxon matched-pairs signed rank test: **P=0.0018) (**i**) and average amplitude (n = 14 cells; Wilcoxon matched-pairs signed rank test: *P=0.0166) of Ca^2+^ events (**j**). (**k, l**) Intensity vs time Ca^2+^ traces from 5 CaMs in control (Ctrl – TTX, 0.5 µM and α1d antagonist BMY7378, 10 µM) (**k**, top) and after PE + BMY application (**l**, top). Heat- map raster plots of the intensity and temporal distribution of Ca^2+^ events in the Ctrl (**k**, bottom) and after PE +BMY application (**l**, bottom). (**m, n**) Graphs showing changes in frequency (n = 14 cells; Paired t-test: *P=0.0303) (**m**) and average amplitude (n = 14 cells; Wilcoxon matched-pairs signed rank test: P=0.0676) of Ca^2+^ events (**n**). All experiments were done in the presence of 0.5 μm TTX. Data are presented as mean ± SEM. ns, not significant. Scale bar, 15 µm.

### NE suppresses proliferation and promotes differentiation of OPCs into oligodendrocytes

Norepinephrine is a potent neuromodulator and can regulate the proliferation and survival of neural progenitors^25, 26^. To study whether NE-mediated Ca^2+^ signaling in OPCs can modulate the fate of these cells, we took a chemogenetic approach to specifically activate LC neurons and induce NE release in the cortex. We cross bred *Dbh-mGC6s* mice with a transgenic mouse line in which hM3Dq, a G_q_ coupled Designer Receptors Exclusively Activated by Designer Drugs (DREADD), and a cytosolic yellow fluorescent protein mCitrine co-expressed under control of a strong ubiquitous CAG promoter in a Cre dependent manner^44^ (*CAG-LSL-HA-hM3Dq-pta-mCitrine;* in short *Gq*) (Fig. 8a). To study the effect of the activation of LC neurons on the fate of cortical OPCs, we treated control (*mGC6s;Gq*) and Dbh-Gq (*Dbh-mGC6s;Gq*) mice for 5 days with CNO (1 mg/kg). Along with CNO, we treated control and *Dbh-Gq* mice with BrdU for two weeks to label proliferating OPCs (Fig 8b). After the end of the BrdU treatment, we performed immunohistochemical analysis using antibodies against eGFP and cell-type specific markers for OPCs and OLs. As expected, the eGFP staining revealed LC fibers in the S1 cortex of *Dbh-Gq* mice, but not in control mice (Fig. 8c, d), confirming that control mice don’t express G_q_ DREADDs in LC neurons. Next, we quantified the number of ASPA+ OLs, PDGFRα+ OPCs and BrdU+ proliferating cells in control and *Dbh-Gq* mice (Fig. 8c-f). This analysis showed a 13.6% increase in the number of ASPA+ OLs (Fig. 8g), and a 38% reduction in the number of BrdU+ cells (Fig. 8h) in *Dbh-Gq* mice in which we exogenously activated NE fibers. However, the total number of PDGFRα+ OPCs remained unchanged between the *control* and *Dbh-Gq* mice (124.6 ± 8.68 and 126.7 ± 15.49, respectively) (Fig. 8i). We did not observe any increase in the number of BrdU+ OLs (<0.5 %), indicating that new OLs were not generated from the newly divided OPCs but from a pre-existing pool of committed OPCs (COPs). The number of BrdU+/PDGFRα+ OPCs was reduced in *Dbh-mGq* (7.4 ± 3.4%) when compared to the control (11.6 ± 4.3%) mice. In summary, results from these experiments support that NE-mediated signaling in OPC can promote their differentiation into OLs and concomitantly suppress their proliferation.

**Figure 8.**
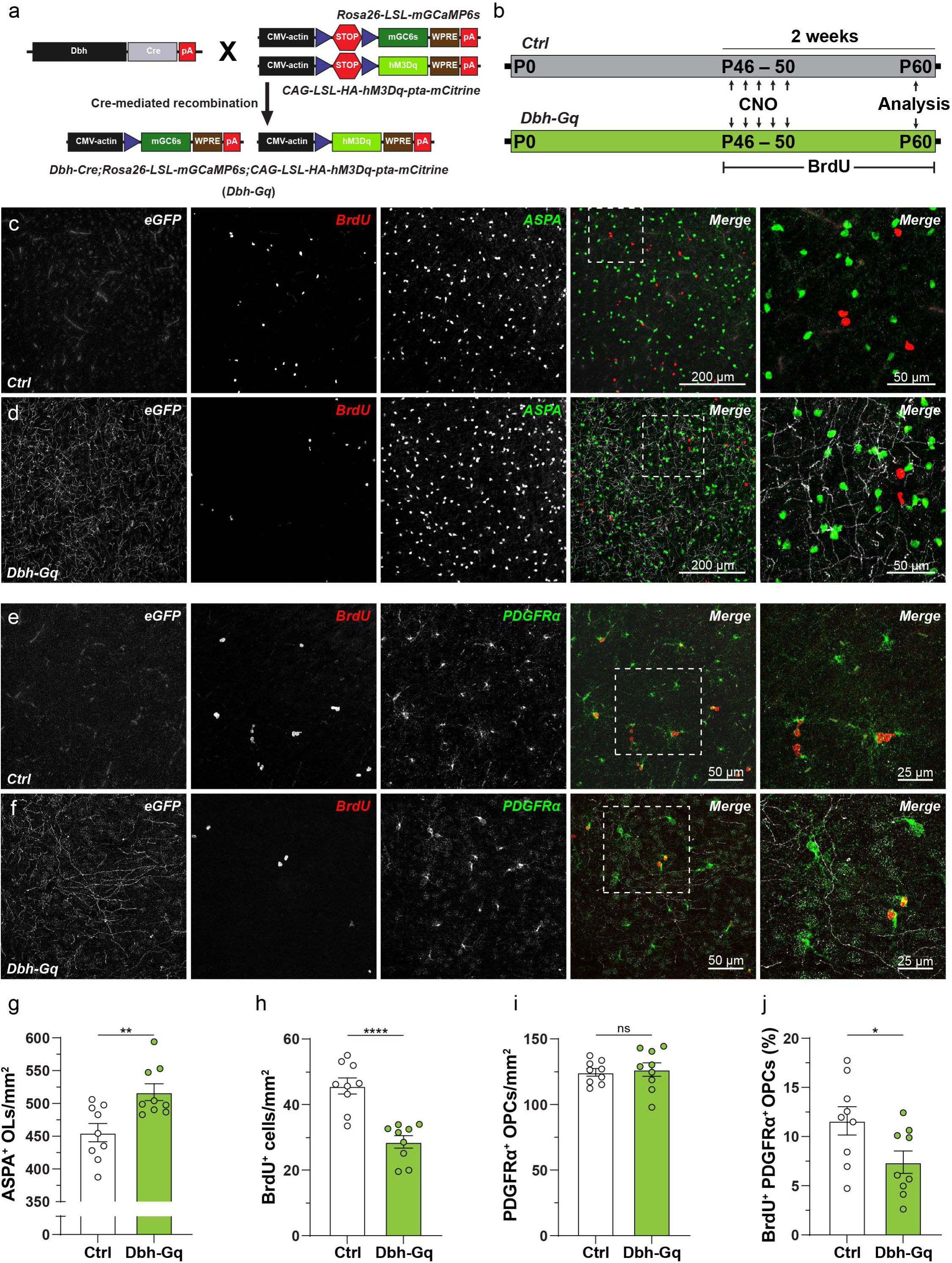
Norepinephrine regulates fate of OPCs by enhancing differentiation and suppressing proliferation. **(a)** Cartoon showing a transgenic strategy to express mGCaMP6s and a chemogenetic effector hM3Dq in noradrenergic (NA) neurons projecting to the cortex in *Rosa26-LSL- mGCaMP6s;CAG-LSL-hM3Dq* (*Control – Ctrl*) and *Dbh-Cre;Rosa26-LSL-mGCaMP6s;CAG-LSL-hM3Dq* (*Dbh-Gq*) mice. (**b**) Strategy to chronically activate NA neurons and induce NE release. 6-7 weeks old *Ctrl* and *Dbh-Gq* mice were injected with CNO for 5 consecutive days to induce NE release, and mice were treated with BrdU for two weeks to label actively diving cells. (**c**, **d**) Maximum intensity projections of confocal z-stacks of the coronal brain sections immunostained stained for eGFP (grey), BrdU (red; cell proliferation marker) and ASPA (green; mature oligodendrocyte marker) from *Ctrl* (**c**) and *Dbh-Gq* (**d**) mice. (right) High-magnification image of the boxed areas in **c, d**. Scale bars, 200 µm and 50 µm (right). (**e, f**) Maximum intensity projections of confocal z-stacks of the coronal brain sections immunostained stained for eGFP (grey), BrdU (red) and PDGFRα (green; OPC marker) from *Ctrl* (**e**) and *Dbh-Gq* (**f**) mice. (right) High-magnification image of the boxed areas in **e, f**. Scale bars, 200 µm and 50 µm (right). (**g**-**i**) Graphs show increase in the number of ASPA^+^ (**g**) (n = 9 slices; Unpaired t-test: **P=0.0049), no changes in the number of PDGFRα^+^ (**h**) (n= 9 slices; Unpaired t-test: P=0.7246) and decreased in the number of BrdU^+^ (**i**) (n = 9 slices; Unpaired t-test: ****P<0.0001) cells in the S1 cortex of control or *Dbh-Gq* mice. (**j**) Graph showing reduced percentage of newly generated PDGFRα^+^ and BrdU^+^ OPCs in *Ctrl* and *Dbh-Gq* mice (n = 9 slices from 3 mice, Unpaired t-test: *P=0.0357). All data are presented as mean ± SEM.

## Discussion

### OPCs exhibit spontaneous and evoked microdomain Ca^2+^ transients

OPCs are small stellate-shaped cells found in both gray and white matter and account for 5- 10% of all neural cells in the adult CNS ranging from fish to mammals. They represent the most abundant group of proliferating cells in the adult CNS. *In vivo* fate mapping studies indicate that they mainly serve as progenitors for OLs during development and continue to do so throughout life. OPCs contribute to the regeneration of OLs and remyelination following experimentally-induced and clinical demyelination. OPCs integrate several signaling pathways including growth factors and neuronal activity before they differentiate into myelinating OLs. Ca^2+^ signaling has been shown to alter the growth and differentiation of cells in a variety of contexts^45^, and Ca^2+^ changes appear to play a prominent role in guiding the fate of OPC^46^. By generating transgenic mice expressing genetically encoded Ca^2+^ sensors in OPC, we discovered that various cell types of oligodendrocyte lineage cells exhibit distinct Ca^2+^ signals, and these signals can be modulated by noradrenaline signaling.

Ca^2+^ transients in OPCs were mainly seen in the processes, and were restricted to a micrometer size local hot-spots of activity (Fig. 2). Such localized hot-spots of cytosolic Ca^2+^ signals, called microdomains, have previously been described in several cell types including neurons^47^ and astrocytes^48^. Somatic Ca^2+^ transients only occurred when mice engaged in intense locomotion activity (Fig. 2), which also led to Ca^2+^ rises across the entire cell. Hence, we suggest that the threshold of summation of CaMs activity in processes to produce cell-wide and somatic Ca^2+^ transients is seemingly quite high in OPCs^49^. The source of microdomain Ca^2+^ fluxes can be both extra- and intracellular and, similar to neurons, CaMs in OPCs might exist in the absence of ultrastructural compartmentalization^47^. However, a detailed electron microscopic analysis will be required to further strengthen this conclusion. About 45% of Ca^2+^ signals in OPCs CaMs were of sub-seconds duration, and ∼1.5 % of all Ca^2+^ events were as fast as 330 ms (Extended Data Fig. 2). It is possible that these sparse but ‘fast’ Ca^2+^ signals represent EPSCs at the axon-OPC synapses generated by the opening of calcium permeable AMPA (cpAMPA), NMDA receptors and A-type voltage gated Ca^2+^ channels in OPCs^36, 49^. Accordingly, in acute brain slices, glutamate and GABA receptor blockers reduced the frequency of Ca^2+^ transients in OPCs, indicating that Ca^2+^ signals in OPCs could be generated by a spontaneous neurotransmitter release. However, the complete block of neuronal firing of action potentials by TTX had no effect on OPCs spontaneous microdomain Ca^2+^ transients (Extended Data Fig. 5). Thus, we propose, as reported in astrocytes, that OPCs have two types of microdomain Ca^2+^ transients – spontaneous and neurotransmitter evoked^37^. Spontaneous Ca^2+^ transients were independent of neuronal activity and could be generated by cell-intrinsic mechanisms such as transient activation of store-operated Ca^2+^ entry, TRP channels^50–52^, reactive oxygen species (ROS), and Ca^2+^ efflux from mitochondria^37, 53, 54^.

### Distinct signatures of microdomain Ca^2+^ signaling in oligodendrocyte lineage cells

Our *in vivo* fate mapping and Ca^2+^ imaging studies revealed that OPCs exhibit distinct Ca^2+^ dynamics while undergoing fateful decision of cell division or differentiation (Fig. 3). Most of the cells with a low frequency of Ca^2+^ events at baseline and little to no increase in Ca^2+^ activity during locomotion had morphological features similar to premyelinating or myelinating OLs (**Group 1 cells** – Fig. 3 and Extended Data Fig. 4). These OLs exhibited Ca^2+^ transients restricted to CaMs as well. OPCs undergo major changes in their physiology when they differentiate into OLs, such as the loss of OPC-axon synapse, the activation of K^+^ leak currents and small fluctuations in membrane potentials^15^. Hence, the source of OLs microdomain Ca^2+^ signals could be more reliant on non-synaptic factors such as activation of TRPA1 channels^50^, opening of mechanotransduction channel Piezo^55^ and G_q_ coupled metabotropic receptors^20^. Also, OLs showed almost no increase in locomotion-induced Ca^2+^ transients, possibly due to downregulation of the expression of adrenergic receptors once the cells had differentiated^20, 56^. With these observations, a key question arises – what is the significance of microdomain Ca^2+^ transients in premyelinating and myelinating OLs? The clue might come from two recent studies on larval zebra fish, which convincingly showed that a modest Ca^2+^ rise in OLs triggers myelin sheath elongation, and high amplitude, long duration Ca^2+^ events induce sheath shortening and retractions events^28, 29^. This indicates that newly generated OLs might integrate information encoded in localized Ca^2+^ transients to trigger myelin formation and fine-tune myelin internodes. However, further detailed analysis will be required to understand the role of OLs Ca^2+^ transients in regulating adult myelination.

Remarkably, some OPCs had characteristics of microdomain Ca^2+^ signals similar to OLs (**Group 1 cells** – Fig. 3). We note that often such OPCs underwent cell-division within few days. In more than 194 imaging sessions, we captured 5 cell-division events. This number matches the previous observations that about 3% of OPCs in the adult brain are in cell-cycle phase^20, 57^. We also found that the summation of the number of CaMs and Ca^2+^ events in the two daughter cells was equal to those in the mother OPCs (Extended Data Fig. 4). Thus, we provide *in vivo* evidence that OPC undergoing cell-division distribute synapses and the repertoire of neurotransmitter receptors equally amongst the two daughter cells – a cell-division feature unique to OPCs^58, 59^. Since NE is known to suppress proliferation of neural progenitors^26^, the non-responsiveness of OPCs to locomotion suggest that OPCs preparing for cell division downregulate expression of adrenergic receptors to relieve them from NE’s brakes on cell-proliferation^27^. We can further support this hypothesis with our observation that chemogenetic activation of LC neurons induced NE release, which suppressed the proliferation of OPCs as well (Fig. 8).

We found a unique class of OPCs that exhibited an exceptionally high baseline frequency of Ca^2+^ events (reaching >100 events/min), but didn’t further increase Ca^2+^ events in response to locomotion activity (**Group 2 cells** – Fig. 3). We suggest that these cells represent a ‘dormant’ pool of OPCs, which might maintain extensive synaptic contacts with unmyelinated axons. Such OPCs might have a high expression level of ionotropic and metabotropic receptors for glutamate and GABA, but low levels of neuromodulator receptors. OPCs can remain in this ‘dormant’ state for several weeks, and are more likely to proliferate than differentiate into OLs^27^ (Fig. 3k). Indeed, we observed that often OPCs shuttled between the dormant and proliferative states (i.e., Group 2 to Group 1 migration). Several studies suggest that activation of AMPA and GABA receptor in OPCs promote their proliferation^60^. It is thus likely that the dormant OPCs integrate neuronal activity at OPC-axon synapses before taking an ultimate decision of cell-division. Accordingly, a recent study combined *in vivo* Ca^2+^ imaging and single-cell RNA sequencing techniques to characterize spinal cord OPCs in larval zebra fish, showed that OPCs with high intracellular Ca^2+^ fluctuations were dormant, and OPCs with low Ca^2+^ fluctuations at baseline tended to differentiate in OLs^27^. Hence, we put forth a hypothesis that the synaptic activity at OPC-axon junctions keep OPCs in a proliferative state^61^, while neuromodulatory signals might guide the fate of OPCs towards death or differentiation.

The most diverse population of oligodendrocyte lineage cells in various transition stages such as OPCs, committed OPCs (COPs) and premyelinating OLs (pMOLs) clustered together in Group 3 (Fig. 3). Unfortunately, due to a lack of distinctive morphological features for OLC cell types, we couldn’t distinguish their precise identity with high certainty in our datasets. In the future, the development of advanced machine learning algorithms to classify cells based on morphological criterion will enable us to identify OPCs, COPs and pMOLs in imaging datasets. Several transcriptomic studies have already classified OLCs into various sub-classes based on their unique mRNA signatures and electrophysiological properties^62^.

One such study described that COPs and pMOLs upregulate the expression of IP_3_R2, which endow these cells with a capacity for an increased responsiveness to the neuromodulator mediated Ca^2+^ signaling^20^. In addition, pMOLs seem to upregulate the expression of α1 adrenergic receptors and become responsive to NE-mediated signaling^20^. We hypothesize that in Group 3, IP_3_R2+ COPs and pMOLs might represent group of cells which can integrate neuromodulatory cues, and are likely to be the ones involved in activity-dependent myelination^20^.

### Activation of adrenergic receptors on OPCs can induce microdomain Ca^2+^ transients

In this study, we uncovered that OPCs show enhanced Ca^2+^ transients in response to the bouts of locomotion during mouse exploratory behavior. Several clues imply that the locomotion induced Ca^2+^ signals in OPCs could be evoked by NE (Fig. 4) – (**1**) It is well established that during an active exploratory phase (i.e. locomotion), when mice are attentive and in the state of vigilance, NE is released by the Dbh^+^ neurons in the locus coeruleus (LC), which project throughout out the brain^39^; (**2**) axonal projection from Dbh^+^ neurons criss-cross the entire cortex and make a close contact with OPC processes; and (**3**) ablation of LC fibers in the cortex, using the neurotoxin DSP4^42^, virtually abolished locomotion-induced activation of CaMs in OPCs. In addition, glutamatergic neurons release neurotransmitters within a few milliseconds once they are depolarized which means that, if glutamate or GABA release at the OPC-axon synapses led to locomotion induced Ca^2+^ transients, OPCs would have responded instantaneously. However, it took about 4 s for OPCs to reach the maximal activation of CaMs from the instant the mouse engaged in locomotion (Fig. 4). Since it takes about 1.87 s for Dbh^+^ neurons to release NE from the time they were depolarized^63^, and the half-life of NE in the brain is about 3.1 s^64^, we infer Ca^2+^ transients in OPCs in response to locomotion are driven by neuromodulators such as NE. Previous studies using transgenic mice expressing eGFP reporters under control of promoters for α1a and α1b ARs showed that α1a ARs are expressed in OPCs but not in mature OLs^56^, and α1b ARs are expressed in both OPCs and OLs^65^. Although α1d-AR promoter-driven LacZ reporter expression shows these receptors are abundant in the cortex, a cell-type specific expression analysis has not been performed^66^. In addition, a recent study on single-cell RNA sequencing of OLCs showed that α1a and α1b ARs were expressed in OLCs, and α1b-ARs was the most expressed sub-type^20^. For a systematic analysis of the functional expression of α1-AR sub-types in OLCs, we performed sm-FISH and extensive α1-ARs pharmacology (Fig. 6 and 7). A detailed sm-FISH analysis with sub-types specific mRNA probes showed that about 45-60% of Olig2+ OLCs expressed various α1-AR sub- types, suggesting that these cells display inherent diversity in the expression of these receptors. Direct activation of α1-ARs on OPCs lead to a rise in intracellular Ca^2+^ concentrations. Hence, we postulate that the state of vigilance and wakefulness in mice induces NE release by LC neurons, which in turn activates α1-ARs on OPCs and induce microdomain Ca^2+^ signals.

### Norepinephrine regulates fate of OPCs and promotes differentiation

NE is known to regulate proliferation and differentiation of neural progenitors^26, 67^. Does NE play a key role in regulating OPCs fate as well? We noted that NE promoted differentiation of OPCs (Fig. 8). However, we also made a peculiar observation - in general, OPC differentiation is coupled to OPC proliferation events^57^. Instead, we observed reduced proliferation of OPCs (Fig. 8). This observation exposed an exciting feature of the effect of NE on the development of the oligodendrocyte lineage. We propose that NE likely didn’t induce differentiation of OPCs themselves, but instead promoted the survival and differentiation of COPs or pMOLs into myelinating OLs. It is well established that pMOLs are susceptible to death and that only a few survive *in vivo*^3, 68^. Although, very little is known about the role of NE signaling on OLCs, reduced levels of NE and hypertrophy of TH^+^ neurons in the LC were reported in a study that carried out a detailed histological analysis on the brains of individuals with multiple sclerosis (MS)^69^. Similar observations were made in the cortex and spinal cord of a mouse model of demyelination (experimental autoimmune encephalomyelitis, EAE), which had reduced NE levels. In addition, studies on such mouse models indicate that increased levels of NE in the CNS can reduce disease severity^69, 70^. Thus, COPs and pMOLs represent a promising pool of cells which effectively respond to NE signaling, and contribute to the adaptive myelination and in myelin repair^71^.

In summary, we have uncovered that OLCs exhibit a wide variety of Ca^2+^ signals, which can be modulated by NE. NE signaling can play a pivotal role in regulating survival, proliferation and differentiation of OLC. Future studies will help us clarify the mechanisms by which this NE signaling contributes to adaptive myelination and myelin plasticity, and explore the therapeutic potential of this mode of signaling to promote remyelination and myelin repair.

## Acknowledgments

We thank Dr. Dwight Bergles (Johns Hopkins University) for generously sharing *Rosa26-lsl- mGCaMP6s*, which were generated by A.A. while he was a post-doctoral fellow in the laboratory of Dr. Bergles. We acknowledge the data storage service SDS@hd supported by the Ministry of Science, Research and the Arts Baden-Württemberg (MWK), and the German Research Foundation (DFG) through grant INST 35/1314–1 FUGG and INST 35/1503–1 FUGG. A.A. was supported in part by the Chica and Heinz Schaller Research Foundation, and grants from the Deutsche Forschungsgemeinschaft: FOR 2289 (P8; AG 287/1-1) and SFB1158 (A09). F.F. is a graduate student the Heidelberg Biosciences International Graduate School (HBIGS), and wants to thank HBIGS for their support.

## Authors contributions

F.F. conducted awake *in vivo* and acute brain slice Ca^2+^ imaging experiments, performed fate mapping studies, contributed to the development of imaging analysis algorithms, and analyzed data. R.R.D characterized transgenic mouse lines used for Ca^2+^ imaging and chemogenetics experiments, performed fate mapping experiments, and sm-FISH based gene expression analysis. K.A. developed and adapted algorithms for analysis of Ca^2+^ imaging data and Mobile HomeCage data. I.C. performed histological characterization of Dbh-Cre mice. A.H. contributed to the characterization of transgenic mice used for Ca^2+^ imaging and acute brain slice Ca^2+^ imaging experiments. A.A. conceived the project, designed the experiments, generated and characterized the *Rosa26-lsl-mGCaMP6s* mice, supervised the project. A.A. and F.F. wrote the manuscript with help from the other authors. All authors have read and approved the manuscript.

## Materials and Methods

Both male and female mice were used for all experiments, and mice were randomly allocated to experimental groups. For *ex vivo e*xperiments, adult mice aged 6–12 weeks old were used and for *in vivo* experiments mice aged 8-16 weeks old were used, unless otherwise described. Mice were maintained on a 12 hours light/dark cycle, and food and water were provided ad libitum. Animal studies were approved by the Governmental Council Karlsruhe, Germany. All animal experiments were carried out in a strict compliance with German Animal Protection Law (TierSCHG) at the Heidelberg University, Germany.

### Generation of ROSA26 targeted conditional mGCaMP6s reporter mouse line

To localize GCaMP6s to the plasma membrane, we fused the gene sequence encoding the first 8 amino acids of the modified MARCKS sequence (MGCCFSKT) to the first methionine (i.e. start ATG) of GCaMP6s sequence (termed mGCaMP6s). To enhance expression of mGCaMP6s, we used a strong ubiquitous CMV-βactin hybrid (CAG) promoter and placed the woodchuck hepatitis virus posttranscriptional regulatory element (WPRE) at the 3′ end of mGCaMP6s expression sequence. For inducible expression of mGCaMP6s, a loxP flanked “stopper” cassette was placed upstream of the mGCaMP76s (Extended Figure 1a). The mGCaMP6s transgenic construct was targeted to the ubiquitously expressed ROSA26 locus using homologous recombination in mouse embryonic stem (ES) cells derived from a SV129 mouse strain. Transgenic mice were generated at the Johns Hopkins University Transgenic Core Laboratory. Germ line transmission was achieved by breeding male chimeric founders to C57Bl/6N wild-type female mice.

### Transgenic Mice

Generation and genotyping of CreER driver lines NG2-CreER^31^ (Jackson Lab 008538), Dbh- Cre^41^, GCaMP6f^72^, tdTomato reporter mouse lines^32^ (Jackson Lab 007914) and hM3Dq- Cirtine chemogenetic effector mice^44^ (Jackson Lab 026220) have been previously described.

### Tamoxifen Injections

The tamoxifen solution for injections (10 mg/mL) was prepared by dissolving tamoxifen (Sigma, T5648) in sunflower seed oil (Sigma, S5007). To ensure the tamoxifen was fully dissolved, the solution was sonicated at room temperature for approximately 5 minutes, then vortex for 30 seconds. This process was repeated until total dissolution was achieved and no crystals remained visible in the solution. The solution was then stored at 4 °C, protected from light, for a maximum of 5 days. Mice aged 4-6 weeks were injected intraperitoneally (i.p.) with tamoxifen at a dosage of 100 mg/kg of body weight once a day for 3 or 5 days, depending on the experiments. Each injection was performed at least 20 hours after the previous one. All subsequent experiments were performed at least 2 weeks after the last tamoxifen injection.

### CNO and BrdU treatment

Clozapine-N-Oxide (CNO) solution was prepared by dissolving water-soluble CNO (HelloBio, HB6149) in 0.9% saline at a concentration of 100 µg/mL. Mice were injected twice with CNO at a dosage of 1.0 mg/kg (i.p.) daily for 5 consecutive days.

During the same period, mice were given 5-Bromo-2’-deoxyuridine (BrdU) (Sigma, B5002). BrdU was dissolved in 0.9% saline at a concentration of 10 mg/mL. Mice were injected with BrdU at a dosage of 50 mg/kg (i.p.) for 5 consecutive days. For maximum labelling of proliferating cells, beginning on the day of the first BrdU injections, mice were also provided with a 1.0 mg/mL BrdU in 1% sucrose solution as drinking water for 14 consecutive days.

### Norepinephrine fiber ablation

To deplete noradrenergic fibers, mice were injected with the noradrenergic neuron specific neurotoxin N-(2-chloroethyl)-N-ethyl-2-bromobenzylamine hydrochloride (DSP4; Sigma, C8417) once at a dose of 50 mg/kg (i.p.). To ensure NE fibers were successfully depleted, subsequent histological and Ca^2+^ imaging experiments were carried out at least 3 days after the DSP-4 injection.

### Immunohistochemistry

Mice were deeply anesthetized by injecting pentobarbital (Narcoren) at 150 mg/kg (i.p.), and transcardially perfused with 4% paraformaldehyde (PFA) in 0.1 M sodium phosphate buffer (pH 7.4). Brains were carefully dissected out from the cranium and post-fixed in 4% PFA for 16 hours (overnight) at 4°C. The brains were then transferred to 1x phosphate-buffered saline (PBS) and 35 µm thick coronal sections were cut using a vibratome (Leica, VT 1000s). Sections were collected in PBS containing 0.2% sodium azide and stored at 4°C until further processed. Prior to immunostaining, the sections were permeabilized 0.3% Triton X-100 in 1x PBS for 10 minutes at room temperature (RT). To prevent non-specific binding of antibodies, the sections were then placed in a blocking buffer (0.3% Triton X-100 and 5% normal donkey serum (NDS) in 1x PBS) for 1 hour at RT. Then, the sections were incubated overnight with the primary antibodies (diluted in blocking buffer) at 4°C with gentle shaking on an orbital shaker. Sections were rinsed 3 times for 5 min each in 1xPBS at RT, and incubated with a blue fluorescent nuclear dye 4′,6-diamidino-2-phenylindole (DAPI) and fluorescent dye-conjugated secondary antibodies (diluted in blocking buffer) for 3 hours at RT. Finally, sections were rinsed thrice for 5 min each in 1x PBS at RT. Sections were then mounted on SuperFrost Plus slides (Thermo Fisher Scientific, 4951PLUS) using Aqua Polymount (Polyscience Inc, 18606-20) and stored at 4°C to dry for at least 16 hours prior to imaging.

Stained coronal sections were then imaged using an epifluorescence microscope (Leica DM6000) and automatically stitched in the accompanying software (LAS X, Leica) upon acquisition. Images were further processed using ImageJ.

### Single RNA-molecule Fluorescent In situ Hybridisation (sm-FISH)

8 weeks old C57BL/6N mice were transcardially perfused with 4% PFA and their brains were isolated and post-fixed as described in the previous section (Immunohistochemistry). Following the post-fixation overnight in 4% PFA at 4°C, the brains were dehydrated by sequentially storing them in 10%, 20% and 30% sucrose in 1xPBS at 4 °C and until they sank to the bottom of the vial in each solution. Brains were then cut to fit a 7 mm x 7 mm × 5 mm (L x W x H) embedding mold, frozen in optimal cutting temperature compound (OCT, Tissue-Tek) with dry ice and stored at −80 °C. The tissue was later sliced into 16 μm thick sections using a cryostat (Leica, CM1950), and stored in an airtight container at −80 °C until further processed. In situ hybridizations was performed using the RNAscope Multiplex Fluorescent Kit v2 (Advanced Cell Diagnostics) for fixed frozen tissue according to the manufacturer’s instructions with the following: Mm-Olig2 (447091), Mm-Adra1a-C2 (408611- C2), Mm-Adra1b-C2 (413561-C2), and Mm-Adra1d (563571). Fluorescent signals were then amplified using a TSA Plus Fluorescein kit (PerkinElmer, NEL741E001KT). In addition, the sections were stained with DAPI and anti-Olig2 antibody (see Immunohistochemistry). All sections were mounted on SuperFrost Plus slides (Thermo Fisher Scientific, 4951PLUS), and later imaged using a confocal microscope (see Confocal Imaging).

smFISH image stacks were imported into Image J for quantification. The nuclei of oligodendrocyte lineage cells were identified using a combination of DAPI and a cell-type specific marker (Olig2), then carefully outlined (dotted line, Fig.6 **a**-**c**). Puncta were manually counted on a plane-by-plane basis from the confocal z-stacks for optimal precision. Cells were only considered positive if the puncta were located directly on the nucleus or its perimeter.

### Confocal Imaging

Images were acquired using a Leica SP8 Confocal microscope equipped with either a 40X (HC PL APO CS2 Water-immersion, Leica) or 63X (HCX PL APO Oil immersion, Leica) objectives, with the pinhole set to 1 Airy Unit (AU). Images were acquired at a resolution of 1024 x 1024 pixels, with a system-optimized step size of 0.423 μm. Image size varied between 290.62 x 290.62 μm and 87.87 x 87,87 μm depending on the objective and magnification factor used. Images were then further processed using ImageJ, and presented as pseudocolored maximum intensity projections.

### Acute Brain Slice Preparation

Mice were anesthetized with isoflurane and immediately decapitated with a pair of large scissors. Their brains were quickly dissected out and stuck on a 5 mm agar stage before being mounted on a vibratome (Leica VT1200S) equipped with a sapphire blade. 250 μm thick coronal cortical slices were cut in ice-cold N-methyl-D-glucamine (NMDG) based cutting solution containing (in mM): 135 NMDG, 1 KCl, 1.2 KH_2_PO_4_, 1.5 MgCl_2_, 0.5 CaCl_2_, 10 Dextrose, 20 Choline Bicarbonate (pH 7.4 and ∼310 mOsm/L). Cortical slices were then incubated in artificial cerebrospinal fluid (ACSF) containing (in mM): 119 NaCl, 2.5 KCl, 2.5 CaCl_2_, 1.3 MgCl_2_, 1 NaH_2_PO_4_, 26.2 NaHCO_3_ and 11 Dextrose (292-298 mOsm/L) for 30 min at 37°C, then maintained at RT for the entire duration of the experiment. The NMDG cutting solution and the ACSF were continuously bubbled with carbogen (95% O_2_/5% CO_2_). All slice imaging experiments were performed at the RT.

### Pharmacological Manipulations

All pharmacological experiments on the acute brain slices were performed by dissolving drugs directly in ACSF, and applying them through a fast perfusion system. Unless otherwise specified, all experiments were performed in the presence of a voltage-gated Na^+^ channel blocker TTX (0.5 μM, HelloBio HB1035) to block neuronal firing of action potentials. Following drugs were used for pharmacological manipulations: adrenergic α1 receptor agonist – phenylephrine (10 μM Sigma P6126); AMPA & Kainate receptors antagonist – CNQX (10 μM, HelloBio HB0205); NMDA receptors antagonist (D-AP5 (50 μM, HelloBio HB0225); Group I & II metabotropic glutamate receptors antagonist – (S)-MCPG (10 μM, HelloBio HB6112); GABA_A_ receptor antagonist – Gabazine (5 μM, HelloBio HB0901); GABA_B_ receptor antagonist – CGP55845 (5 μM; HelloBio HB0960); adrenergic α1_A_ receptor antagonist – RS17053 (40 μM, Tocris 0985); adrenergic α1_B_ receptor antagonist – Chloroethylclonidine (30 μM, Sigma-Aldrich B003); and adrenergic α1_D_ receptors antagonist – BMY 7378 (10 μM, Tocris 1006).

### Time-Lapse Two-Photon Microscopy in Acute Brain Slices

Intracellular Ca^2+^ concentration changes and morphology of OLCs in the acute cortical brain slices were imaged on a custom-built 2-photon microscope (Bergamo II, ThorLabs) fitted with a 20X water-immersion objective (1.0NA, XLUMPLFLN, Olympus), 8 kHz galvo/resonance scanner, piezo drive for fast z-scanning, and two GaAsP PMT detectors. 2-photon excitation for GCaMP6f, mGCaMP6s and tdTomato at a single wavelength was achieved using a mode locked Ti:Sapphire pulsed laser (Chameleon Ultra II, Coherent) tuned at 940 nm. Images were acquired at a resolution of 512 x 512 pixels (85,5 x 85,5 μm) and 12 bit pixel depth, with each imaging session consisting of a 5-minute recording of 920 frames at 3.1 Hz. Over the entire course of the imaging slices were steadily perfused with oxygenated ASCF. Since it was difficult to identify cells expressing mGCaMP6s, slices were briefly exposed to 10µM phenylephrine prior to each experiment, and responsive cells suitable for further imaging were identified.

### Glass Cranial Windows Implantation

Mice aged 5-8 weeks old were implanted with chronic glass cranial windows^37, 73^. Mice were first anesthetized using isoflurane (5% for induction, 1.5%-2% during surgery, mixed with 0.5L/min O_2_), and placed on a custom stereotaxic frame (RWD) fitted with a stereo microscope (Stemi 508, Zeiss) and thermostat-controlled heating pad (RWD). Throughout the surgery mice were maintained at 37 °C body temperature, and were administered with dexamethasone (2 mg/kg, i.p.) and 0.03 mg/kg buprenorphine sub-cutaneously (s.c.) to reduce inflammation and pain, respectively. Skin on the scalp was removed to expose the skull over the right cerebral hemisphere, where a 2 x 2 mm area (centered at 1 mm below Bregma and 3 mm lateral from the midline) was marked using a scalpel. This perimeter was gently drilled to remove the bone, and 2 x 2 mm piece of cover glass (#1, Menzel-Gläser) was placed within the craniotomy and sealed to the skull using histoacrylamine glue (Histoacryl, Braun). A helicopter headbar with a 5.2 mm circular opening (model 2, NeuroTar) was attached using dental cement (Kulzer, Paladur) mixed with cyanoacryl (Hager Werken, 152261). After the surgery, mice were administered carprofen (5 mg/kg, s.c.) every 12 hours for two days to reduce pain and inflammation. Whenever possible, multiple mice were housed in the same cage with 1-2 wooden shelters, providing them with an enriched environment. Mice were allowed to recover from surgery for two weeks, before Mobile HomeCage training could begin.

### Mobile Home Cage Setup and Training

All intravital two-photon imaging sessions in awake animals were performed using a high precision 2D locomotion tracking device (Mobile HomeCage (MHC), NeuroTar). The MHC consists of a solid aluminum platform coupled to an air compressor which allows a constant upwards flow of air through holes on its surface. The uniform air-cushion it creates is sufficient to support an ultralight carbon-fiber cage (180 mm diameter, 70 mm height) with negligible friction. When mice were placed inside, they were able to easily displace the carbon-fiber cage and engage in unrestricted self-motivated exploration while still being head-fixed, enabling us to study Ca^2+^ dynamics in OLCs and mouse behavior simultaneously. In addition, high-precision magnetic trackers embedded in the MHC platform constantly calculate the position and the speed of the animal inside the experimental setup using two small magnets located at the bottom of the carbon cage.

To ensure mice are comfortable being head-fixed in the MHC, mice were progressively acclimatized over the course of one week to remain head-fixed for about 60 min during awake 2-photon microscopy sessions in the MHC. On the first two days, the mice were introduced to the 2-photon imaging environment and extensively handled by the experimenter. The handling steps included occasional manual head-restriction to simulate imaging conditions inside the MHC. On the third day, mice were placed for ∼15 minutes inside the carbon-fiber cage of MHC floated on the air-cushion, and could freely explore the imaging environment without head-fixation. Finally, over the next 4 days, the mice were head-fixed in MHC and placed under the 2-photon microscope for an increasing period of time (+ 10 minutes per day), replicating the exact conditions of an imaging session. By the end of the 7^th^ day of habituation, mice were fully habituated and minimally stressed while head-fixated in MHC, and were ready for imaging session lasting between 45-60 min.

### Intravital Two-Photon Microscopy on Awake Behaving Mice

As previously mentioned, intravital 2-photon microscopy in awake mice was performed using the MHC setup, enabling us to correlate Ca^2+^ dynamics in OLCs with explorative behavior of mice. To ensure synchronization of behavioral and Ca^2+^ imaging data, image acquisition on the 2-photon microscope was initiated by a TTL pulse from the Mobile HomeCage motion tracking software (Tracker version 2.1.10.3). Images were acquired on a custom-built 2- photon microscope (Bergamo II, ThorLabs) with a 20X water-immersion objective (1.0NA, XLUMPLFLN, Olympus). 2-photon excitation was achieved using a mode locked Ti:Sapphire pulsed laser (Chameleon Ultra II, Coherent) tuned at 940 nm (see Time-Lapse Two-Photon Microscopy in Acute Brain Slices section for more details). Unless otherwise specified, image time-series were acquired at a rate of 5,1 Hz and a resolution of 512 x 512 pixels (85,5 x 85,5 μm) for a total of 10 minutes. Mice were kept head fixed on MHC for a maximum of one hour per imaging day.

### Mobile HomeCage Data Analysis

All data about the location and speed of mice in the Mobile HomeCage were imported to MATLAB (Mathworks, 2021a) using a custom code. Active periods were detected automatically and aligned with the 2-photon imaging data using a custom MATLAB code (Mathworks, 2021a). Locomotion bouts were defined as the period during which the speed of the mouse was above threshold, which was set at one standard deviation above the mean speed of the animal during that session. Importantly, some mice displayed low levels of motor activity, significantly lowering the mean speed and standard deviation used to establish the threshold, which could lead to “pseudo-bouts” bouts being detected. Hence, bouts were excluded if they did not contain speeds above 20% of that of the maximum observed in that session. In addition, to account for residual locomotion-induced Ca^2+^ activity that occurs immediately after the end of an active phase, each locomotion bout was extended by 2.41 seconds, covering about 92% of interspike intervals observed during locomotion. Mice were considered to be in a resting state when the speed remained below threshold for an uninterrupted period of at least 30 s. Ca^2+^ signals in OLCs were sorted into “Resting” or “Active” categories corresponding to the type of bout during which these events were initiated. Infrequent short locomotion bouts devoid of any Ca^2+^ signals were excluded from any further analysis.

### Preprocessing of Image Stack for Ca^2+^ Signal Analysis

To correct for movement artefacts in XY plane, 2-photon time-series image stacks were post hoc registered using automatic alignment algorithm TurboReg^74^ (plugin for ImageJ). During registration TurboReg transforms images into a 32-bit format. Therefore, after registration, images were reformatted to 16-bit. Surrounding background pixels that were not part of the cell (based on median-intensity projection of the image stack) including pericytes and larger blood vessels were cropped to accelerate analysis. For an automatic analysis of Ca^2+^ microdomain and Ca^2+^ signals in OLCs, our machine learning based algorithm CaSCaDe was modified accordingly for specific imaging datasets (see details below).

### Extraction and Analysis of Ca^2+^ signal in Oligodendrocyte Lineage cells and (5 Hz)

Using the average projection of the 2-photon image stack, we automatically generated a Ca^2+^ microdomain map as described in Agarwal et al. 2017^37^. Prior to event detection, the traces from each Ca^2+^ microdomain were extracted and flattened using the MATLAB “detrend” function. Since awake in vivo imaging can lead to movement-induced changes in fluorescence that are independent of the Ca^2+^ concentration, we also extracted fluorescence signals from the static tdTomato channel using the same Ca^2+^ microdomain map. Because the fluorescence of tdTomato does not change as a function of Ca^2+^ concentration, any rapid changes in fluorescence observed in this channel can be interpreted as an artifact. To avoid including such artifacts in our analysis, we thus subtracted the tdTomato traces from those of GCaM6f for microdomains. Ca^2+^ events were automatically detected using peak detection algorithm of CaSCaDe. Only Ca^2+^ events with amplitude more than 5 z-scores and duration at least 4 frames were considered. For each Ca^2+^ event, the beginning and ending points were determined using the point of maxima for each peak as a reference. Beginning of the peak was defined as the last point preceding the maximum that has a value below 1 z-score, and the ending is defined as a point where 1 z-score is reached for two consecutive frames, or falls within 10% of the starting point’s value. The resulting Ca^2+^ signals were classified based on a trained SVM classifier^37^.

### Fast imaging (15 Hz)

To acquire image stack at a 15 Hz sampling frequency, the imaging was performed at lower frame averaging rate (@2) and resolution (256 x 256 pixels), which resulted in higher pixel sizes (0.17 μm/pixel at 5 Hz vs 0.33 μm/pixel at 15 Hz) and increased background noise.

Hence, CaSCaDe was modified as follows to accommodate images with lower signal to noise ratio. First, a filtering step was added to exclude inactive or highly noisy microdomains that did not exhibit clear (7+ z-score) Ca^2+^ events. Next, peak detection was performed on the selected microdomains as described in the previous section. Since images acquired at 15 Hz are inherently noisy, accurately estimating the duration of Ca^2+^ events was not possible using the methods implemented to analyze images acquired at 5Hz. To resolve this, the microdomain Ca^2+^ traces were passed through a 1 Hz lowpass filter. Then, start and end points of Ca^2+^ events were computed separately for both the original unfiltered and filtered traces. This method allowed for optimal detection of both fast events (using the unfiltered traces) and slow events (using the filtered traces) simultaneously. Hence, a combination of both binary detection signals (unfiltered and filtered) was computed to obtain the final duration of each event. Importantly, hyper-segmented events (i.e. events that were detected separately in the unfiltered trace, but that correspond to a single event in the filtered trace) were combined if they occurred less than 0.2 seconds apart. The resulting Ca^2+^ signals were also classified based on a trained SVM classifier^37^.

### Noradrenergic fibers

Due to high noise levels, the automatic segmentation on CaSCaDe resulted in spurious microdomain maps with hypersegmentation of ROIs. Using the average projection of the imaging stacks, noradrenergic fibers were manually identified and regions of interest (ROIs) delineated. Manual ROIs were imported into MATLAB as binary masks and used as a template for CaSCaDe to generate the Ca^2+^ microdomain map and to detect Ca^2+^ events.

### Statistical Analysis

Statistical analyses were performed using GraphPad Prism (version 9.4.0). Each dataset was individually tested to assess the normality of its distribution using the D’Agostino-Pearson normality test. If both datasets passed the normality test, a two-tailed t-test was used to determine the statistical significance of the difference between two groups. Otherwise, a non-parametric test (Wilcoxon-matched pairs signed rank test) was used for paired comparisons. If three or more datasets were compared, we used Kruskall-Wallis test with Dunn’s correction to verify statistical significance between groups. The tests used for each comparison (and their significance level) are indicated in the figure legends. Data are shown in the figures as mean ± SEM and p<0.05 was considered significant. *N* represents the number of animals and *n* the number of cells (physiology) or brains slices (histology).

## Tables

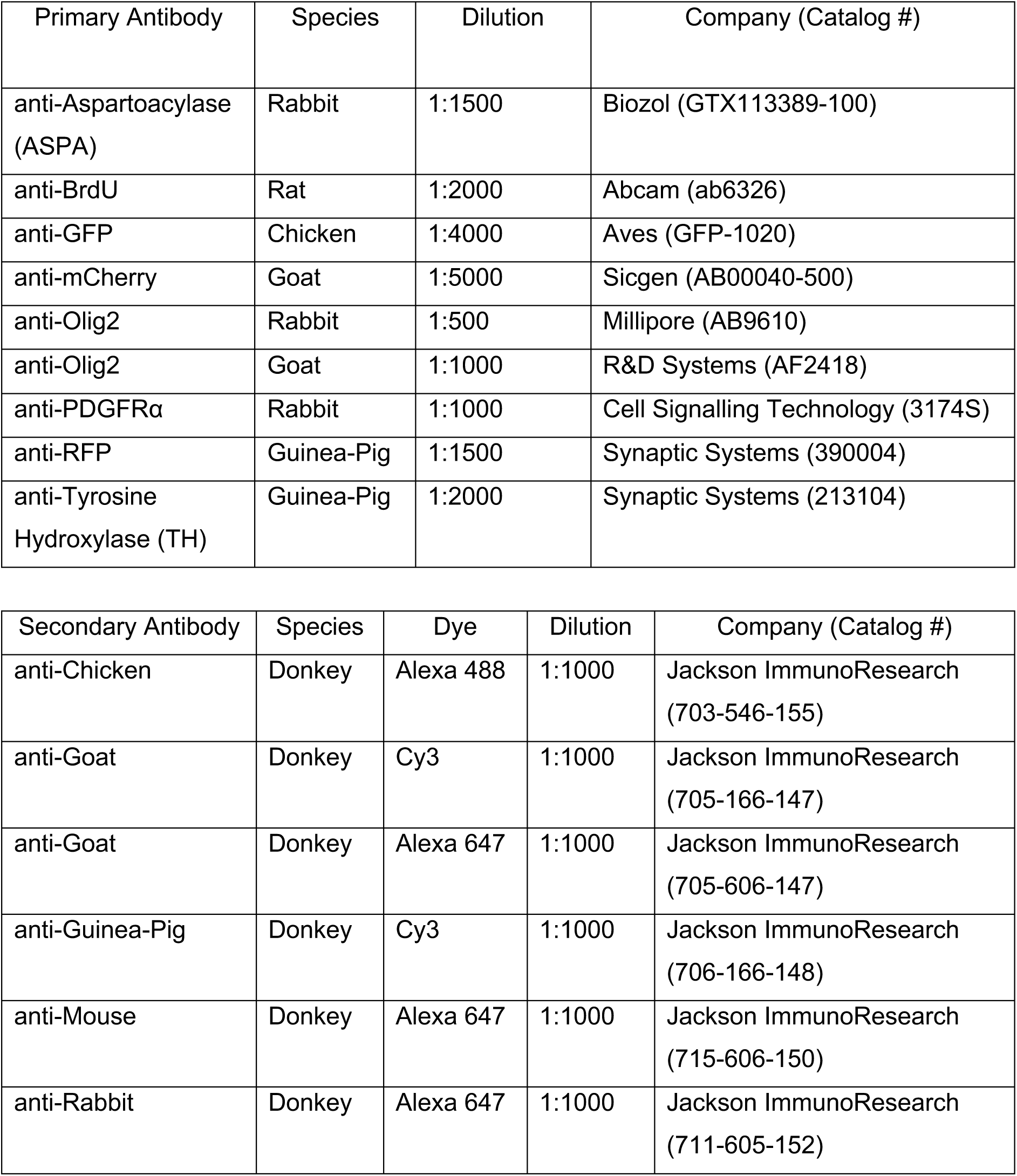
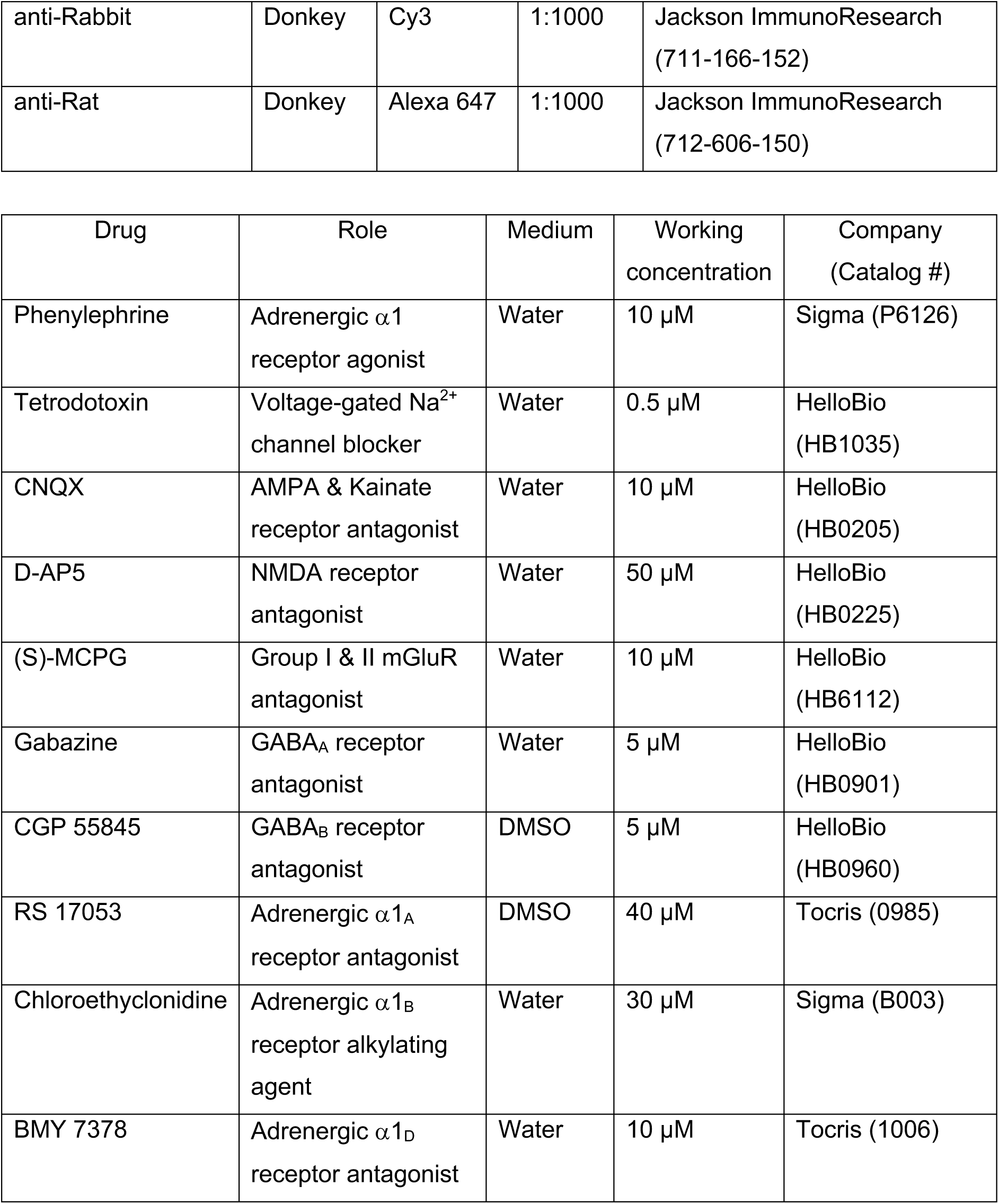

| Primary Antibody | Species | Dilution | Company (Catalog #) |
| --- | --- | --- | --- |
| anti-Aspartoacylase (ASPA) | Rabbit | 1:1500 | Biozol (GTX113389-100) |
| anti-BrdU | Rat | 1:2000 | Abcam (ab6326) |
| anti-GFP | Chicken | 1:4000 | Aves (GFP-1020) |
| anti-mCherry | Goat | 1:5000 | Sicgen (AB00040-500) |
| anti-Olig2 | Rabbit | 1:500 | Millipore (AB9610) |
| anti-Olig2 | Goat | 1:1000 | R&D Systems (AF2418) |
| anti-PDGFRα | Rabbit | 1:1000 | Cell Signalling Technology (3174S) |
| anti-RFP | Guinea-Pig | 1:1500 | Synaptic Systems (390004) |
| anti-Tyrosine Hydroxylase (TH) | Guinea-Pig | 1:2000 | Synaptic Systems (213104) |

| Secondary Antibody | Species | Dye | Dilution | Company (Catalog #) |
| --- | --- | --- | --- | --- |
| anti-Chicken | Donkey | Alexa 488 | 1:1000 | Jackson ImmunoResearch (703-546-155) |
| anti-Goat | Donkey | Cy3 | 1:1000 | Jackson ImmunoResearch (705-166-147) |
| anti-Goat | Donkey | Alexa 647 | 1:1000 | Jackson ImmunoResearch (705-606-147) |
| anti-Guinea-Pig | Donkey | Cy3 | 1:1000 | Jackson ImmunoResearch (706-166-148) |
| anti-Mouse | Donkey | Alexa 647 | 1:1000 | Jackson ImmunoResearch (715-606-150) |
| anti-Rabbit | Donkey | Alexa 647 | 1:1000 | Jackson ImmunoResearch (711-605-152) |
| anti-Rabbit | Donkey | Cy3 | 1:1000 | Jackson ImmunoResearch<br>(711-166-152) |
| anti-Rat | Donkey | Alexa 647 | 1:1000 | Jackson ImmunoResearch<br>(712-606-150) |

| Drug | Role | Medium | Working concentration | Company (Catalog #) |
| --- | --- | --- | --- | --- |
| Phenylephrine | Adrenergic $\alpha_1$ receptor agonist | Water | 10 μM | Sigma (P6126) |
| Tetrodotoxin | Voltage-gated Na<sup>2+</sup> channel blocker | Water | 0.5 μM | HelloBio (HB1035) |
| CNQX | AMPA & Kainate receptor antagonist | Water | 10 μM | HelloBio (HB0205) |
| D-AP5 | NMDA receptor antagonist | Water | 50 μM | HelloBio (HB0225) |
| (S)-MCPG | Group I & II mGluR antagonist | Water | 10 μM | HelloBio (HB6112) |
| Gabazine | GABA<sub>A</sub> receptor antagonist | Water | 5 μM | HelloBio (HB0901) |
| CGP 55845 | GABA<sub>B</sub> receptor antagonist | DMSO | 5 μM | HelloBio (HB0960) |
| RS 17053 | Adrenergic $\alpha_{1A}$ receptor antagonist | DMSO | 40 μM | Tocris (0985) |
| Chloroethylclonidine | Adrenergic $\alpha_{1B}$ receptor alkylating agent | Water | 30 μM | Sigma (B003) |
| BMY 7378 | Adrenergic $\alpha_{1D}$ receptor antagonist | Water | 10 μM | Tocris (1006) |

---

**Extended Data Figure 1.**
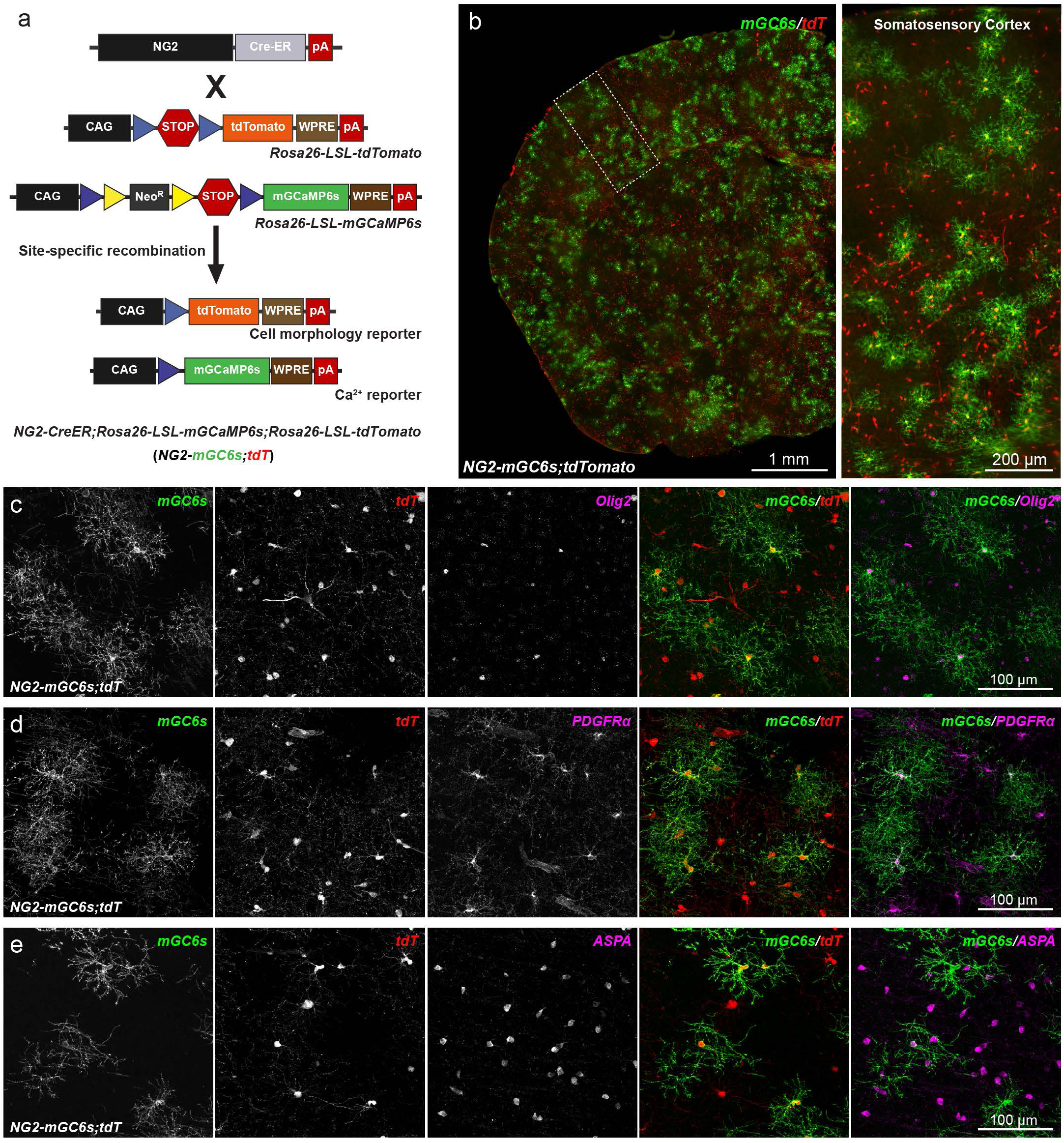
Conditional expression of mGCaMP6s and tdTomato in oligodendrocyte lineage cells. (**a**) Cartoon showing a triple transgenic strategy to express membrane anchored variant of GCaMP6s (mGCaMP6s) and tdTomato in OPCs using *NG2-CreER;Rosa26-LSL- mGCaMP6s;Rosa26-LSL-tdTomato* (*NG2-mGC6s;tdT*) mice. (**b**) (left) Coronal hemi-section of brain from *NG2-mGC6s;tdT* mouse stained for mGCaMP6s (mGC6s, green) and tdTomato (tdT, red). (right) Zoom-in image from the S1 cortex (boxed area shown in the left panel) showing oligodendrocyte lineage cells co-expressing mGC6s and tdT. Scale bars, 1 mm (left) and 200 µm (right). (**c**) Confocal microscopy images showing double recombined (mGC6s^+^/tdT^+^) cells expressing pan oligodendrocyte lineage marker Olig2 (magenta). (**d**) Images showing a subset mGC6s^+^/tdT^+^ cells were PDGFRα^+^ OPCs (in magenta). (**e**) Images showing a subset of recombined mGC6s^+^/tdT^+^ cells expressing ASPA (magenta), a marker for mature oligodendrocytes. Scale bars, 100 µm (**c-e**).

**Extended Data Figure 2.**
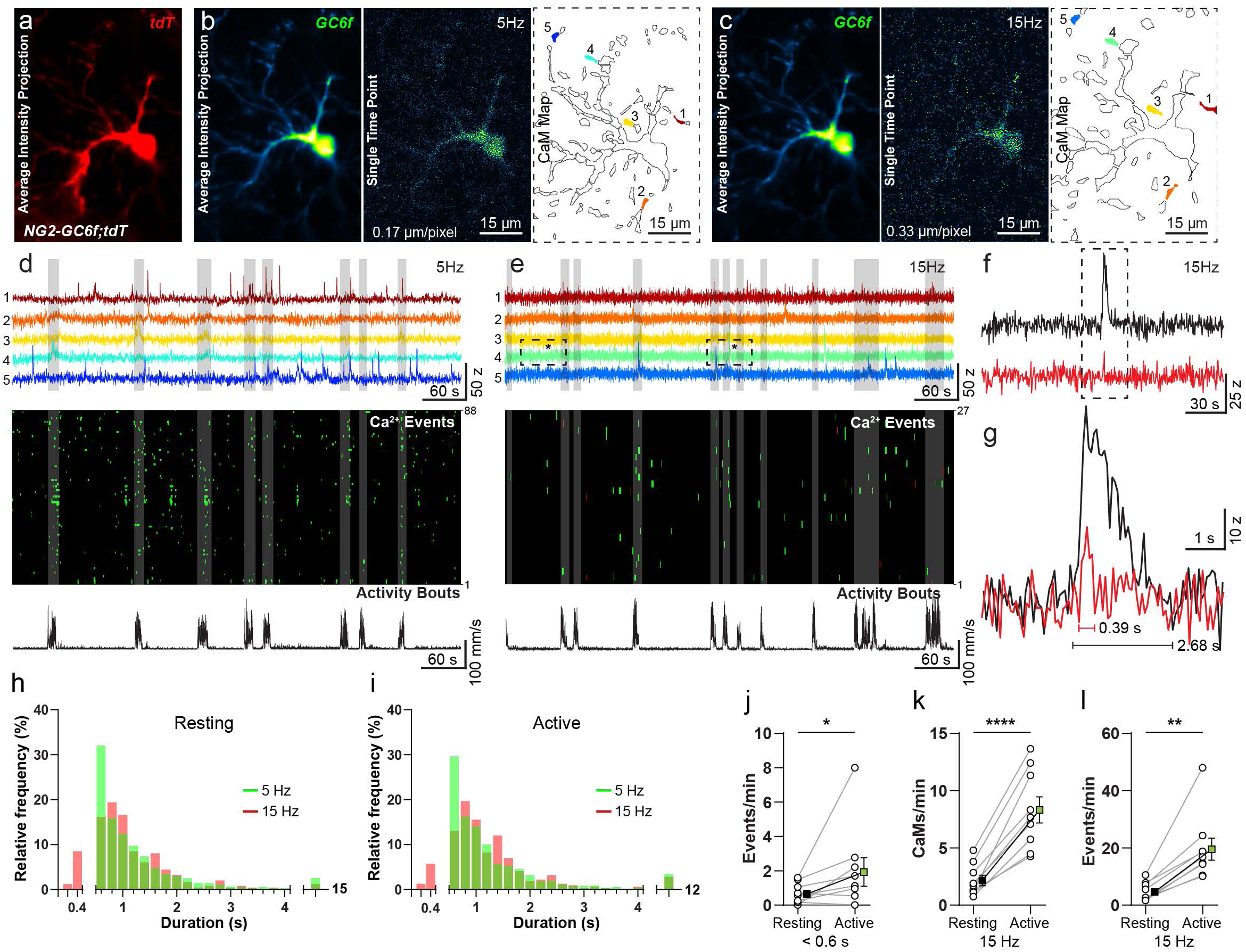
OPCs exhibit low frequency but fast Ca^2+^ transients *in vivo* in awake behaving mice. (**a, b**) Average intensity time-series projection of tdTomato expressed by an OPC in the S1 cortex of *NG2-GC6f;tdT* mouse. (**b**) Average intensity time-series projection of GCaMP6f (left), a single time frame (center), and map of 88 detected CaMs (right) of a 10-minute image stack acquired at 5 Hz. (**c**) Average intensity time-series projection of GCaMP6f (left), a single time frame (center), and map of 27 detected Ca^2+^ microdomains (right) of a 10- minute image stack acquired at 15 Hz. (**d, e**) Intensity vs time Ca^2+^ traces from 5 CaMs (top) at 5 Hz (**d**) and 15 Hz (**e**) corresponding to colors in **b**, **c.** Binarized raster plots of the temporal distribution of Ca^2+^ events (middle) at 5Hz (**d**) and 15 Hz (**e**). Fast Ca^2+^ events with duration < 600ms only detectable at 15Hz are shown in red (**e,** middle). Mouse locomotion activity trace (bottom) at 5Hz (**d**) and 15 Hz (**e**). Vertical grey bars in **d, e** indicate bouts of mouse locomotive activity. (**f**) Magnified Ca^2+^ signal peaks from the boxed areas in green trace (**e**). (**g**) Zoom-in of a fast (red) and slow (black) Ca^2+^ events detected using 15 Hz imaging speed, and their respective durations. (**h**, **I**) Frequency distribution histograms of duration of Ca^2+^ events detected at 5 Hz (green) and 15 Hz (red) during resting phase (**h**) and active locomotion bouts (**i**). The first segment of the x-axis shows the distribution of Ca^2+^ events that are only detected at 15 Hz acquisition rate. Last segment of x-axis shows distribution of Ca^2+^ events >4 s duration. (**j)**, Graph showing changes in the frequency of ‘fast’ Ca^2+^ events (<600ms) detected only at an acquisition rate of 15 Hz (n = 9 cells; Wilcoxon matched-pairs signed rank test: *P=0.0391). (**k**, **l**) Graphs showing increase in the number of Ca^2+^ microdomains (**k**) (n = 9 cells; Paired t-test: ****P<0.0001) and frequency of Ca^2+^ events (**l**) (n = 9 cells; Wilcoxon matched-pairs signed rank test: **P=0.0039) during resting and active phases at 15 Hz acquisition rate. All data are presented as mean ± SEM. Scale bar, 15 µm.

**Extended Data Figure 3.**
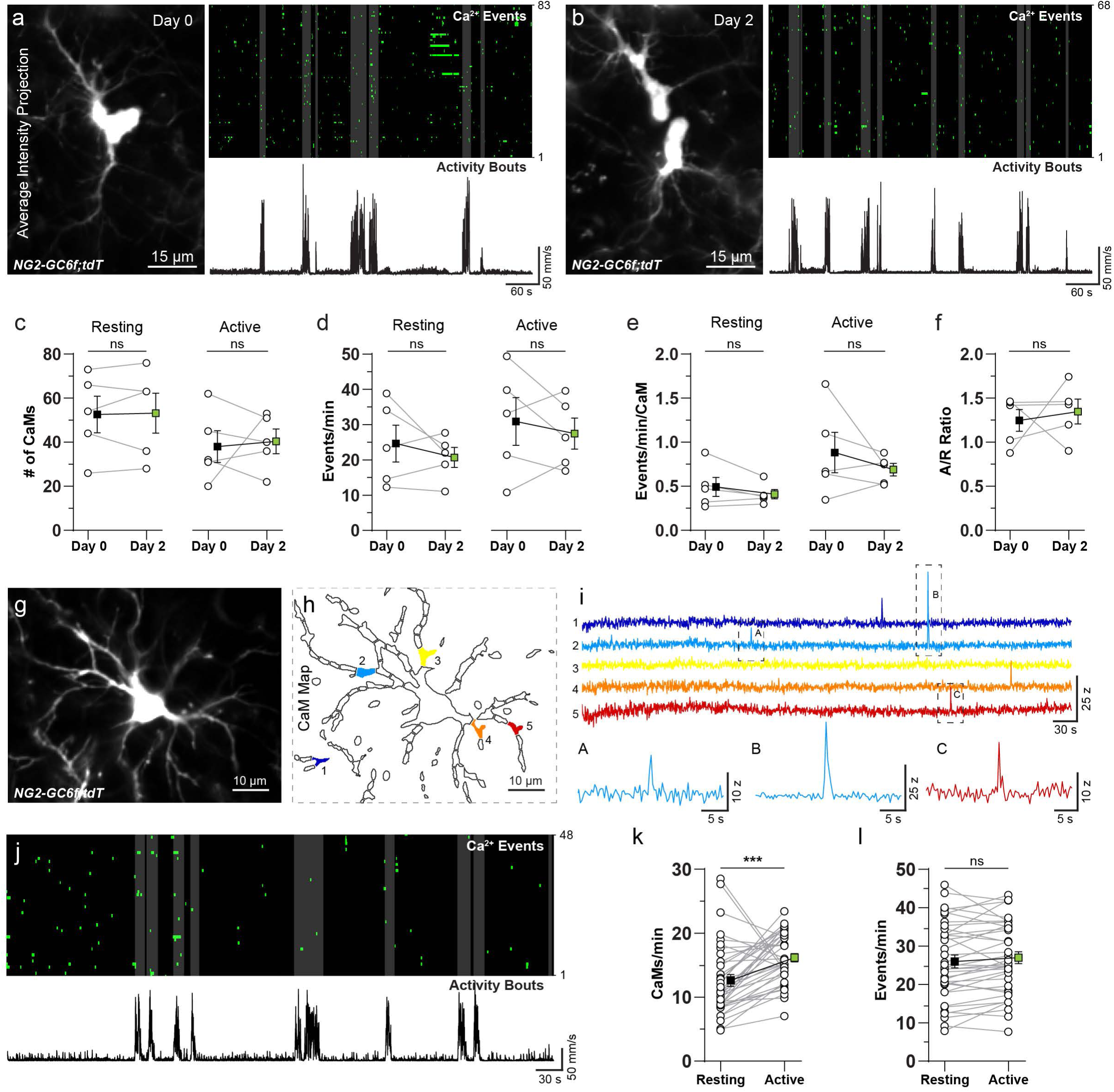
Characteristics of Ca^2+^ signals in OPCs undergoing cell division and differentiation into oligodendrocyte. (**a**) (left) Average intensity time-series projection of tdTomato reporter expressed by an OPC in the S1 cortex of *NG2-GC6f;tdT* mouse before (**a,** Day 0) and after (**b,** Day 2) cell division event. (right, top) Binarized raster plot show Ca^2+^ events occurred in 83 CaMs (Day 0) and 68 CaMs (Day 2) during a 10-min imaging period, and (right, bottom) mouse locomotion activity trace. Vertical grey bars in the raster plot represent bouts of locomotion. (**c**-**e**) Graphs comparing the number of active Ca^2+^ microdomains (**c**), frequency of Ca^2+^ events (**d**), the frequency of Ca^2+^ events per Ca^2+^ microdomain (**e**) during resting phase (left) and active locomotion bouts (right) before and after cell division event (n = 5 cells; Wilcoxon matched-pairs signed rank test: P>0.05). (**f**) Graph comparing the ratio between the number of Ca^2+^ events per minute detected during active and resting phases, before (Day 0, left) and after (Day 2, right) cell division (n = 5 cells; Wilcoxon matched-pairs signed rank test: P>0.05). (**g, h**) Average intensity time-series projection of tdTomato reporter expressed by a differentiated oligodendrocyte in the S1 cortex of *NG2-GC6f;tdT* mouse (**g**) and its associated CaM map (**h**). (**i**) (top) Intensity vs time Ca^2+^ traces from 5 CaMs corresponding to colors in **h**. (bottom) Magnified Ca^2+^ signal peaks from the boxed areas (A, B, C). (**j**) Binarized raster plot showing detected Ca^2+^ events in all 48 CaMs (top) occurring during a 10-min imaging period, and mouse locomotion activity trace (bottom). Vertical grey bars in the raster plot represent bouts of locomotion. (**k**, **l**) Graphs comparing number of Ca^2+^ microdomains (**k**) (n = 37 cells; Wilcoxon matched-pairs signed rank test: ***P=0.0003), and frequency of Ca^2+^ events (**l**) (n = 37 cells; Paired t-test: P=0.1257; ns, not significant) in differentiated oligodendrocyte during the resting and active locomotion phases. All data are presented as mean ± SEM. Scale bars: 15 µm (**a, b**) and 10 μm (**g**).

**Extended Data Figure 4.**
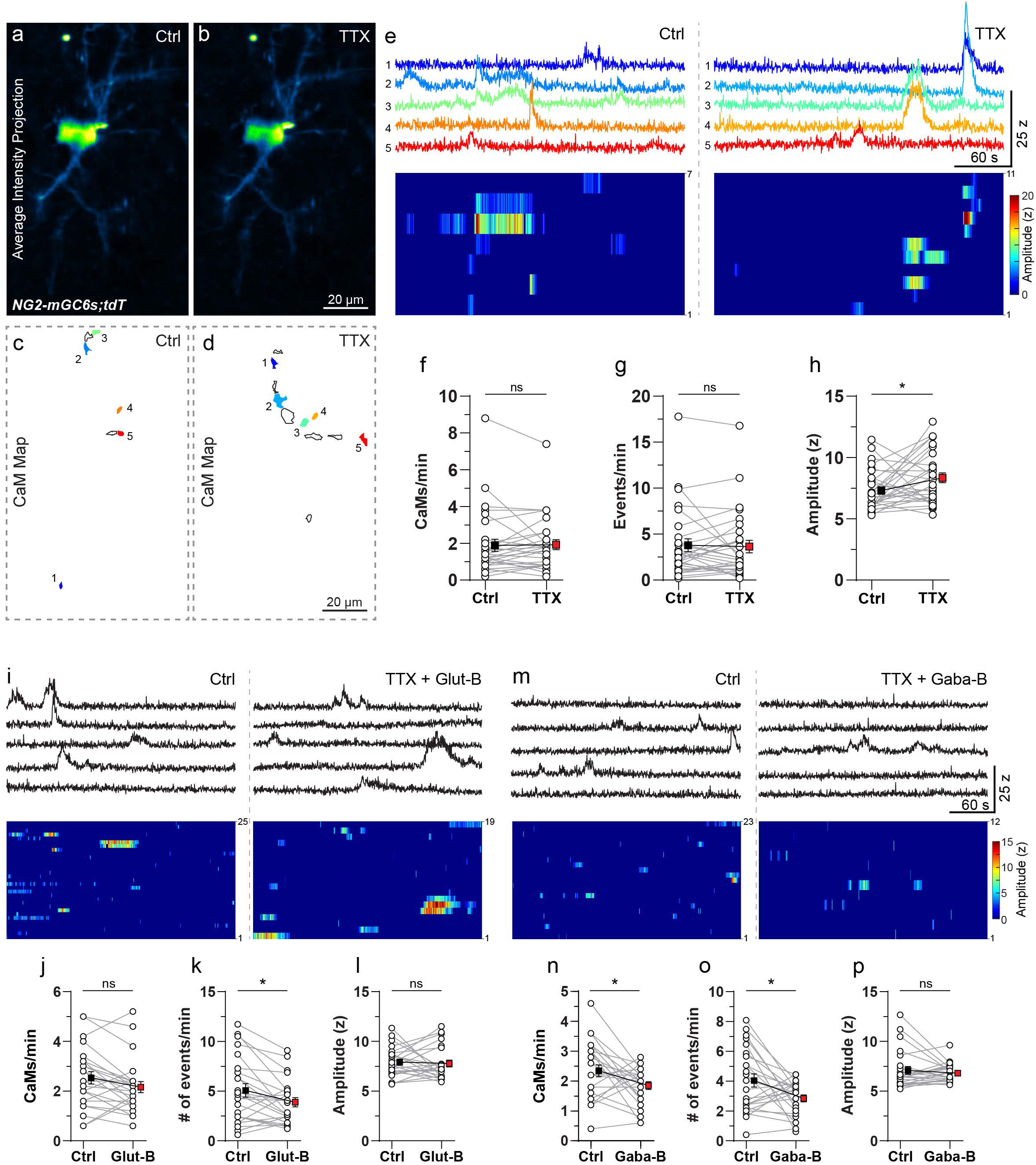
OPCs exhibit spontaneous Ca^2+^ transients which persist in the presence of neuronal activity blockers. (**a**, **b**) Average intensity time-series projection of mGCaMP6s signal in an OPC in an acute brain slice from *NG2-mGC6s;tdT* mouse in control (Ctrl) (**a**) and after bath application of voltage gated sodium channel blocker TTX (0.5 µM) (**b**). (**c, d**) CaM maps recorded in Ctrl conditions (**c**) and after TTX application (**d**). (**e**) Intensity vs time Ca^2+^ traces from 5 CaMs in Ctrl (top, left) and after TTX application (top, right) corresponding to colors in **c, b**. Heat-map raster plots of the intensity and temporal distribution of Ca^2+^ events in the Ctrl (bottom, left) and after TTX application (bottom, right). (**f**-**h**) Graphs comparing number of active Ca^2+^ microdomains per minute (**f**) (n = 29 cells; Wilcoxon matched-pairs signed rank test: P=0.3394), frequency of Ca^2+^ events (**g**) (n = 29 cells; Wilcoxon matched-pairs signed rank test: P=0.9777), and the amplitude of Ca^2+^ events (**h**) (n = 29 cells; Paired t-test: *P=0.0365) in Ctrl conditions and after TTX application. (**i**) Intensity vs time Ca^2+^ traces from 5 CaMs in Ctrl (top, left) and after glutamate receptor blockers (Glut-B: TTX 0.5µM; CNQX, 10 µM; AP5,50 µM; MCPG, 10 µM) application (top, right). Heat-map raster plots of the intensity and temporal distribution of Ca^2+^ events in the Ctrl (bottom, left) and after Glut-B application (bottom, right). (**j-l**) Graphs comparing the number of Ca^2+^ microdomains per minute (**j**) (n = 24 cells; Wilcoxon matched-pairs signed rank test: P=0.0666), the frequency of Ca^2+^ events (**k**) (n = 24 cells; Paired t-test: *P=0.014), and the amplitude of Ca^2+^ events (**l**) (n = 24 cells; Paired t-test: P=0.7268) in Ctrl and after Glut-B application. (**m**) Intensity vs time Ca^2+^ traces from 5 CaMs in Ctrl (top, left) and after GABA receptor blockers (Gaba-B: TTX 0.5 µM; CGP 55845, 5 µM; Gabazine, 5 µM; left) application (top, right). Heat-map raster plots of the intensity and temporal distribution of Ca^2+^ events in the Ctrl (bottom, left) and after Gaba-B application (bottom, right). (**n-p**) Graphs comparing the number of Ca^2+^ microdomains per minute (**o**) (n = 24 cells; Paired t-test: *P=0.0369), frequency of Ca^2+^ events (**n**) (n = 24 cells; Paired t-test: *P=0.0182), and amplitude of Ca^2+^ events (**p**) (n = 24 cells; Wilcoxon matched-pairs signed rank test: P=0.8996) in control and after Gaba-B application. All data are presented as mean ± SEM. ns, not significant. Scale bars, 20 µm.

**Supplementary Video 1. OPC Ca^2+^ transients in the S1 cortex of awake, freely-moving mice engaging in explorative behavior.** Pseudocolored time-series images of GCaMP6f in a cortical OPC (top left) and its dynamic Ca^2+^ microdomain (CaM) map overlaid on the average intensity projection of the static reporter (tdTomato) (top right). (bottom) 2D representations of the position of the mouse in the MHC (left, grey circle), its speed (center), and the number of active CaMs at any given time. Images were acquired at 5.1 frames/s and displayed at 75 frames/s. Scale bar, 10 µm.

**Supplementary Video 2. Ca^2+^ transients of divided OPCs in the S1 cortex of awake, freely-moving mice.** (top left) Pseudocolored time-series images of GCaMP6f in splitting cortical OPCs. (top right) Dynamic Ca^2+^ microdomain map of daughter cell 1 (red, left) and daughter cell 2 (green, right) overlaid on the average intensity projection of the static reporter (tdTomato). (bottom) Graphs of the locomotion speed of the mouse in the MHC (left), and the number of simultaneously active CaMs in daughter cell 1 (red, center) and daughter cell 2 (green, right) at any given time. Images were acquired at 5.1 frames/s and displayed at 75 frames/s. Scale bar, 10 µm.

**Supplementary Video 3. Ca^2+^ transients of an OL in the S1 cortex of awake, freely-moving mice.** Pseudocolored time-series images of GCaMP6f in a cortical OL (top left) and its dynamic Ca^2+^ microdomain (CaM) map overlaid on the average intensity projection of the static reporter (tdTomato) (top right). (bottom) 2D representations of the position of the mouse in the MHC (left, grey circle), its speed (center), and the number of simultaneously active CaMs at any given time. Images were acquired at 5.1 frames/s and displayed at 75 frames/s. Scale bar, 10 µm.

**Supplementary Video 4. Ca^2+^ transients of noradrenergic fibers in the S1 cortex of awake, freely-moving mice.** Pseudocolored time-series images of mGCaMP6s in noradrenergic fibers in cortex (top left) and its dynamic Ca^2+^ microdomain (CaM) map overlaid on the average time-series projection of mGCaMP6s signal (top right). (bottom) 2D representations of the position of the mouse in the MHC (left, grey circle), its speed (center), and the number of simultaneously active CaMs at any time. Images were acquired at 5.1 frames/s and displayed at 75 frames/s. Scale bar, 10 µm.

**Supplementary Video 5. Spontaneous and phenylephrine-evoked Ca^2+^ transients in a cortical OPC from an acute brain slice.** Pseudocolored time-series images of mGCaMP6s in a cortical OPC (left) with its dynamic Ca^2+^ microdomain (CaM) map overlaid on the average intensity projection of the static reporter (tdTomato) (right). Images were acquired at 3.1 frames/s and displayed at 75 frames/s. Time (0 - 277s): spontaneous Ca^2+^ events (in TTX, 0.5 µM); Time (278 - 550s): phenylephrine evoked Ca^2+^ events (TTX, 0.5 µM + PE, 10 µM). Scale bar, 10 µm.

## References

1. Tripathi, R. B. et al. Remarkable Stability of Myelinating Oligodendrocytes in Mice. Cell Reports 21, 316–323 (2017).

2. Yeung, M. S. Y. et al. Dynamics of Oligodendrocyte Generation and Myelination in the Human Brain. Cell 159, 766–774 (2014).

3. Hughes, E. G., Orthmann-Murphy, J. L., Langseth, A. J. & Bergles, D. E. Myelin remodeling through experience-dependent oligodendrogenesis in the adult somatosensory cortex. Nat Neurosci 21, 696–706 (2018).

4. Seggie, J. & Berry, M. Ontogeny of interhemispheric evoked potentials in the rat: Significance of myelination of the corpus callosum. Exp Neurol 35, 215–232 (1972).

5. Tomassy, G. S. et al. Distinct Profiles of Myelin Distribution Along Single Axons of Pyramidal Neurons in the Neocortex. Science 344, 319–324 (2014).

6. Nave, K.-A. Myelination and support of axonal integrity by glia. Nature 468, 244–252 (2010).

7. Fields, R. D. Myelination: An Overlooked Mechanism of Synaptic Plasticity? Neurosci 11, 528–531 (2005).

8. Paez, P. M. & Lyons, D. A. Calcium Signaling in the Oligodendrocyte Lineage: Regulators and Consequences. Annu Rev Neurosci 43, 1–24 (2020).

9. Xin, W. & Chan, J. R. Myelin plasticity: sculpting circuits in learning and memory. Nat Rev Neurosci 21, 682–694 (2020).

10. Stadelmann, C., Timmler, S., Barrantes-Freer, A. & Simons, M. Myelin in the Central Nervous System: Structure, Function, and Pathology. Physiol Rev 99, 1381–1431 (2019).

11. Franklin, R. J. M. & ffrench-Constant, C. Remyelination in the CNS: from biology to therapy. Nat Rev Neurosci 9, 839–855 (2008).

12. Lundgaard, I. et al. Neuregulin and BDNF Induce a Switch to NMDA Receptor-Dependent Myelination by Oligodendrocytes. Plos Biol 11, e1001743 (2013).

13. Ziskin, J. L., Nishiyama, A., Rubio, M., Fukaya, M. & Bergles, D. E. Vesicular release of glutamate from unmyelinated axons in white matter. Nat Neurosci 10, 321–330 (2007).

14. Kukley, M., Capetillo-Zarate, E. & Dietrich, D. Vesicular glutamate release from axons in white matter. Nat Neurosci 10, 311–320 (2007).

15. Larson, V. A., Zhang, Y. & Bergles, D. E. Electrophysiological properties of NG2+ cells: Matching physiological studies with gene expression profiles. Brain Res 1638, 138–160 (2016).

16. Kukley, M., Nishiyama, A. & Dietrich, D. The Fate of Synaptic Input to NG2 Glial Cells: Neurons Specifically Downregulate Transmitter Release onto Differentiating Oligodendroglial Cells. J Neurosci 30, 8320–8331 (2010).

17. Biase, L. M. D., Nishiyama, A. & Bergles, D. E. Excitability and Synaptic Communication within the Oligodendrocyte Lineage. J Neurosci 30, 3600–3611 (2010).

18. Goebbels, S. et al. A neuronal PI(3,4,5)P3-dependent program of oligodendrocyte precursor recruitment and myelination. Nat Neurosci 20, 10–15 (2017).

19. Lee, S.-H. & Dan, Y. Neuromodulation of Brain States. Neuron 76, 209–222 (2012).

20. Marques, S. et al. Oligodendrocyte heterogeneity in the mouse juvenile and adult central nervous system. Science 352, 1326–1329 (2016).

21. Zhang, Y. et al. An RNA-Sequencing Transcriptome and Splicing Database of Glia, Neurons, and Vascular Cells of the Cerebral Cortex. J Neurosci 34, 11929–11947 (2014).

22. Abiraman, K. et al. Anti-Muscarinic Adjunct Therapy Accelerates Functional Human Oligodendrocyte Repair. J Neurosci 35, 3676–3688 (2015).

23. Mei, F. et al. Micropillar arrays as a high-throughput screening platform for therapeutics in multiple sclerosis. Nat Med 20, 954–960 (2014).

24. Berridge, C. W. & Waterhouse, B. D. The locus coeruleus–noradrenergic system: modulation of behavioral state and state-dependent cognitive processes. Brain Res Rev 42, 33–84 (2003).

25. Kulkarni, V. A., Jha, S. & Vaidya, V. A. Depletion of norepinephrine decreases the proliferation, but does not influence the survival and differentiation, of granule cell progenitors in the adult rat hippocampus. Eur J Neurosci 16, 2008–2012 (2002).

26. Weselek, G. et al. Norepinephrine is a negative regulator of the adult periventricular neural stem cell niche. Stem Cells 38, 1188–1201 (2020).

27. Marisca, R. et al. Functionally distinct subgroups of oligodendrocyte precursor cells integrate neural activity and execute myelin formation. Nat Neurosci 23, 363–374 (2020).

28. Krasnow, A. M., Ford, M. C., Valdivia, L. E., Wilson, S. W. & Attwell, D. Regulation of developing myelin sheath elongation by oligodendrocyte calcium transients in vivo. Nat Neurosci 21, 24–28 (2018).

29. Baraban, M., Koudelka, S. & Lyons, D. A. Ca2+ activity signatures of myelin sheath formation and growth in vivo. Nat Neurosci 21, 19–23 (2018).

30. Daigle, T. L. et al. A Suite of Transgenic Driver and Reporter Mouse Lines with Enhanced Brain-Cell-Type Targeting and Functionality. Cell 174, 465–480.e22 (2018).

31. Zhu, X. et al. Age-dependent fate and lineage restriction of single NG2 cells. Development 138, 745–753 (2011).

32. Madisen, L. et al. A robust and high-throughput Cre reporting and characterization system for the whole mouse brain. Nat Neurosci 13, 133–140 (2010).

33. Benediktsson, A. M., Schachtele, S. J., Green, S. H. & Dailey, M. E. Ballistic labeling and dynamic imaging of astrocytes in organotypic hippocampal slice cultures. J Neurosci Meth 141, 41–53 (2005).

34. Broussard, G. J. et al. In vivo measurement of afferent activity with axon-specific calcium imaging. Nat Neurosci 21, 1272–1280 (2018).

35. Kislin, M. et al. Flat-floored Air-lifted Platform: A New Method for Combining Behavior with Microscopy or Electrophysiology on Awake Freely Moving Rodents. J Vis Exp Jove 51869 (2014) doi:10.3791/51869.

36. Chen, T.-W. et al. Ultrasensitive fluorescent proteins for imaging neuronal activity. Nature 499, 295–300 (2013).

37. Agarwal, A. et al. Transient Opening of the Mitochondrial Permeability Transition Pore Induces Microdomain Calcium Transients in Astrocyte Processes. Neuron 93, 587–605.e7 (2017).

38. Wei, Z. et al. A comparison of neuronal population dynamics measured with calcium imaging and electrophysiology. Plos Comput Biol 16, e1008198 (2020).

39. Paukert, M. et al. Norepinephrine Controls Astroglial Responsiveness to Local Circuit Activity. Neuron 82, 1263–1270 (2014).

40. Ye, L. et al. Ethanol abolishes vigilance-dependent astroglia network activation in mice by inhibiting norepinephrine release. Nat Commun 11, 6157 (2020).

41. Parlato, R., Otto, C., Begus, Y., Stotz, S. & Schütz, G. Specific ablation of the transcription factor CREB in sympathetic neurons surprisingly protects against developmentally regulated apoptosis. Development 134, 1663–1670 (2007).

42. Ross, S. B. & Stenfors, C. DSP4, a Selective Neurotoxin for the Locus Coeruleus Noradrenergic System. A Review of Its Mode of Action. Neurotox Res 27, 15–30 (2015).

43. Perez, D. M. α1-Adrenergic Receptors in Neurotransmission, Synaptic Plasticity, and Cognition. Front Pharmacol 11, 581098 (2020).

44. Zhu, H. et al. Cre-dependent DREADD (Designer Receptors Exclusively Activated by Designer Drugs) mice. Genesis 54, 439–446 (2016).

45. Clapham, D. E. Calcium Signaling. Cell 131, 1047–1058 (2007).

46. Butt, A. M. Neurotransmitter-mediated calcium signalling in oligodendrocyte physiology and pathology. Glia 54, 666–675 (2006).

47. Goldberg, J. H., Tamas, G., Aronov, D. & Yuste, R. Calcium Microdomains in Aspiny Dendrites. Neuron 40, 807–821 (2003).

48. Grosche, J. et al. Microdomains for neuron–glia interaction: parallel fiber signaling to Bergmann glial cells. Nat Neurosci 2, 139–143 (1999).

49. Sun, W., Matthews, E. A., Nicolas, V., Schoch, S. & Dietrich, D. NG2 glial cells integrate synaptic input in global and dendritic calcium signals. Elife 5, e16262 (2016).

50. Hamilton, N. B., Kolodziejczyk, K., Kougioumtzidou, E. & Attwell, D. Proton-gated Ca2+- permeable TRP channels damage myelin in conditions mimicking ischaemia. Nature 529, 523–527 (2016).

51. Shigetomi, E., Jackson-Weaver, O., Huckstepp, R. T., O’Dell, T. J. & Khakh, B. S. TRPA1 Channels Are Regulators of Astrocyte Basal Calcium Levels and Long-Term Potentiation via Constitutive d-Serine Release. J Neurosci 33, 10143–10153 (2013).

52. Shigetomi, E., Tong, X., Kwan, K. Y., Corey, D. P. & Khakh, B. S. TRPA1 channels regulate astrocyte resting calcium and inhibitory synapse efficacy through GAT-3. Nat Neurosci 15, 70–80 (2012).

53. Battefeld, A., Popovic, M. A., Vries, S. I. de & Kole, M. H. P. High-Frequency Microdomain Ca2+ Transients and Waves during Early Myelin Internode Remodeling. Cell Reports 26, 182–191.e5 (2019).

54. Rui, Y., Pollitt, S. L., Myers, K. R., Feng, Y. & Zheng, J. Q. Spontaneous Local Calcium Transients Regulate Oligodendrocyte Development in Culture through Store-Operated Ca2+ Entry and Release. Eneuro 7, ENEURO.0347-19.2020 (2020).

55. Coste, B. et al. Piezo proteins are pore-forming subunits of mechanically activated channels. Nature 483, 176–181 (2012).

56. Cohen, R. I. & Almazan, G. Norepinephrine-stimulated PI hydrolysis in oligodendrocytes is mediated by alpha 1A-adrenoceptors. Neuroreport 4, 1115–8 (1993).

57. Hughes, E. G., Kang, S. H., Fukaya, M. & Bergles, D. E. Oligodendrocyte progenitors balance growth with self-repulsion to achieve homeostasis in the adult brain. Nat Neurosci 16, 668–676 (2013).

58. Ge, W.-P., Zhou, W., Luo, Q., Jan, L. Y. & Jan, Y. N. Dividing glial cells maintain differentiated properties including complex morphology and functional synapses. Proc National Acad Sci 106, 328–333 (2009).

59. Kukley, M. et al. Glial cells are born with synapses. Faseb J 22, 2957–2969 (2008).

60. Khawaja, R. R. et al. GluA2 overexpression in oligodendrocyte progenitors promotes postinjury oligodendrocyte regeneration. Cell Reports 35, 109147 (2021).

61. Chen, T.-J. et al. In Vivo Regulation of Oligodendrocyte Precursor Cell Proliferation and Differentiation by the AMPA-Receptor Subunit GluA2. Cell Reports 25, 852–861.e7 (2018).

62. Spitzer, S. O. et al. Oligodendrocyte Progenitor Cells Become Regionally Diverse and Heterogeneous with Age. Neuron 101, 459–471.e5 (2019).

63. Huang, H.-P. et al. Long latency of evoked quantal transmitter release from somata of locus coeruleus neurons in rat pontine slices. Proc National Acad Sci 104, 1401–1406 (2007).

64. Palij, P. & Stamford, J. A. Real-time monitoring of endogenous noradrenaline release in rat brain slices using fast cyclic voltammetry: 1. Characterisation of evoked noradrenaline efflux and uptake from nerve terminals in the bed nucleus of stria terminalis, pars ventralis. Brain Res 587, 137–146 (1992).

65. Papay, R. et al. Mouse α1B-adrenergic receptor is expressed in neurons and NG2 oligodendrocytes. J Comp Neurol 478, 1–10 (2004).

66. Sadalge, A. et al. α1d Adrenoceptor signaling is required for stimulus induced locomotor activity. Mol Psychiatr 8, 664–672 (2003).

67. Bortolotto, V. et al. Salmeterol, a β2 Adrenergic Agonist, Promotes Adult Hippocampal Neurogenesis in a Region-Specific Manner. Front Pharmacol 10, 1000 (2019).

68. Trapp, B. D., Nishiyama, A., Cheng, D. & Macklin, W. Differentiation and Death of Premyelinating Oligodendrocytes in Developing Rodent Brain. J Cell Biology 137, 459–468 (1997).

69. Polak, P. E., Kalinin, S. & Feinstein, D. L. Locus coeruleus damage and noradrenaline reductions in multiple sclerosis and experimental autoimmune encephalomyelitis. Brain 134, 665–677 (2011).

70. Simonini, M. V. et al. Increasing CNS Noradrenaline Reduces EAE Severity. J Neuroimmune Pharm 5, 252–259 (2010).

71. Hughes, E. G. & Stockton, M. E. Premyelinating Oligodendrocytes: Mechanisms Underlying Cell Survival and Integration. Frontiers Cell Dev Biology 9, 714169 (2021).

72. Madisen, L. et al. Transgenic Mice for Intersectional Targeting of Neural Sensors and Effectors with High Specificity and Performance. Neuron 85, 942–958 (2015).

73. Holtmaat, A. et al. Long-term, high-resolution imaging in the mouse neocortex through a chronic cranial window. Nat Protoc 4, 1128–1144 (2009).

74. Thevenaz, P., Ruttimann, U. E. & Unser, M. A pyramid approach to subpixel registration based on intensity. Ieee T Image Process 7, 27–41 (1998).

